# Tracing Endometriosis: Coupling deeply phenotyped, single-cell based Endometrial Differences and AI for disease pathology and prediction

**DOI:** 10.1101/2024.08.09.606959

**Authors:** Lea Duempelmann, Shaoline Sheppard, Angelo Duo, Jitka Skrabalova, Brett McKinnon, Thomas Andrieu, Dennis Goehlsdorf, Sukalp Muzumdar, Cinzia Donato, Ryan Lusby, Wiebke Solass, Hans Bösmüller, Peter Nestorov, Michael D. Mueller

## Abstract

Endometriosis, affecting 1 in 9 women, presents treatment and diagnostic challenges. To address these issues, we generated the biggest single-cell atlas of endometrial tissue to date, comprising 466,371 cells from 35 endometriosis and 25 non-endometriosis patients without exogenous hormonal treatment. Detailed analysis reveals significant gene expression changes and altered receptor-ligand interactions present in the endometrium of endometriosis patients, including increased inflammation, adhesion, proliferation, cell survival, and angiogenesis in various cell types. These alterations may enhance endometriosis lesion formation and offer novel therapeutic targets. Using ScaiVision, we developed neural network models predicting endometriosis of varying disease severity (median AUC = 0.83), including an 11-gene signature-based model (median AUC = 0.83) for hypothesis-generation without external validation. In conclusion, our findings illuminate numerous pathway and ligand-receptor changes in the endometrium of endometriosis patients, offering insights into pathophysiology, targets for novel treatments, and diagnostic models for enhanced outcomes in endometriosis management.

## Introduction

Endometriosis is a prevalent gynecological disease affecting approximately 11% of women of reproductive age worldwide^1^ and imposing substantial societal and economic burdens due to its chronic nature and debilitating symptoms^2^. Characterized by the abnormal growth of endometrial tissue outside the uterus, endometriosis manifests through chronic pelvic pain, intense pain during periods and intercourse, and infertility^3,4^. The diagnosis of endometriosis is challenging due to the variability in clinical symptoms and the absence of confirmed in vitro diagnostics (IVDs)^5^. This results in a 6 to 11-year delay between the onset of symptoms and diagnosis with surgical laparoscopy^6^. The delay not only prolongs patients’ suffering from inadequate management strategies but also lets the untreated disease advance to more severe stages^7^.

Current limited treatment options and diagnostic challenges underscore the need for a comprehensive understanding of the pathophysiology and accurate in vitro diagnostic tools for endometriosis. Sampson’s widely accepted 1927 theory postulated the migration of shed endometrial tissue during menstruation to the peritoneal cavity, where it forms endometriotic lesions^8^. Since most women experience retrograde menstruation, other risk factors have been suggested to contribute to the development of endometriosis, such as differences in the endometrium^9,10^.

Monthly hormonal influences drive extensive endometrial remodeling, encompassing proliferation, lineage specialization, and vascularization, resulting in marked functional and morphological changes that render the endometrium an exceptionally complex tissue to study. The largest endometrium bulk-sequencing studies with over 200 patients and stratified by menstrual cycle phases reported no significant gene expression changes between women with and without endometriosis^11,12^. Nonetheless, bulk sequencing may mask influential gene expression changes of rare yet critical cell types in such highly complex tissue. The advent of single-cell sequencing has revolutionized our ability to dissect complex tissues at an unprecedented resolution^13^. However, recent single-cell endometriosis studies including the eutopic endometrium faced limitations such as small eutopic sample size, cell type exclusion, exogenous hormone treatment, or did not consider menstrual cycle phase changes^14–19^. A refined single-cell study of entire endometrium biopsies from non-hormonally treated patients throughout the menstrual cycle is currently lacking.

In this study, we present the largest single-cell atlas of endometrial tissue to date, comprising 466,371 cells from 35 endometriosis and 25 control patients without exogenous hormones at different menstrual cycle phases. Through detailed cell annotation, we identify significant gene expression alterations in various endometrial cell types among patients with endometriosis, particularly in pathways related to inflammation, adhesion, proliferation, cell survival, and angiogenesis. Furthermore, several receptor-ligand interactions between various cell types were altered, pointing toward a pivotal role for macrophages and stimulated inflammation, angiogenesis, proliferation, and cell survival, which may favor endometriosis susceptibility. Leveraging convolutional neural networks through ScaiVision, we have developed models that can effectively predict mild and severe endometriosis with high accuracy based on either all endometrial cells or exclusively cycling mesenchymal cells. Additionally, we have successfully distilled a concise signature comprising only 11 genes, which retains remarkable accuracy in predicting endometriosis. In conclusion, the findings from our single-cell atlas reveal several altered pathways and ligand-receptor interactions in different cell types, offering new targets for more effective non-hormonal treatments, and have led to the generation of precise in vitro diagnostic models for endometriosis.

## Results

### Generation of the cycle stage annotated single-cell endometrial atlas for the investigation of endometriosis

To confidently characterize cells within the endometrium, identify transcriptomic changes across the menstrual cycle, and reveal endometriosis-specific differences, we performed single-cell sequencing (10x Genomics) on endometrial biopsies collected from 60 women prior to laparoscopic surgery. Of these women, 35 were confirmed to have endometriosis through histopathological evaluation (ENDO) and 25 patients were endometriosis-free (non-ENDO) (Fig. 1a). The menstrual cycle phase of each patient was assessed from serum progesterone levels, histological endometrium evaluation (if available) by two independent pathologists, and the transcriptional profile. Samples from different menstrual cycle phases were included, as were mild (ASRM stages I-II, mild-ENDO) and severe (ASRM stages III-IV, severe-ENDO) endometriosis cases (Fig. 1b). Any patients taking exogenous hormones within 3 months before surgery were excluded, due to the diverse impact of hormones on the transcriptional landscape of the endometrium^20^ (Extended Data Fig. 3g,h). Additional exclusion criteria encompassed other inflammatory diseases, malignancy, pregnancy, and lactation.

**Fig. 1.**
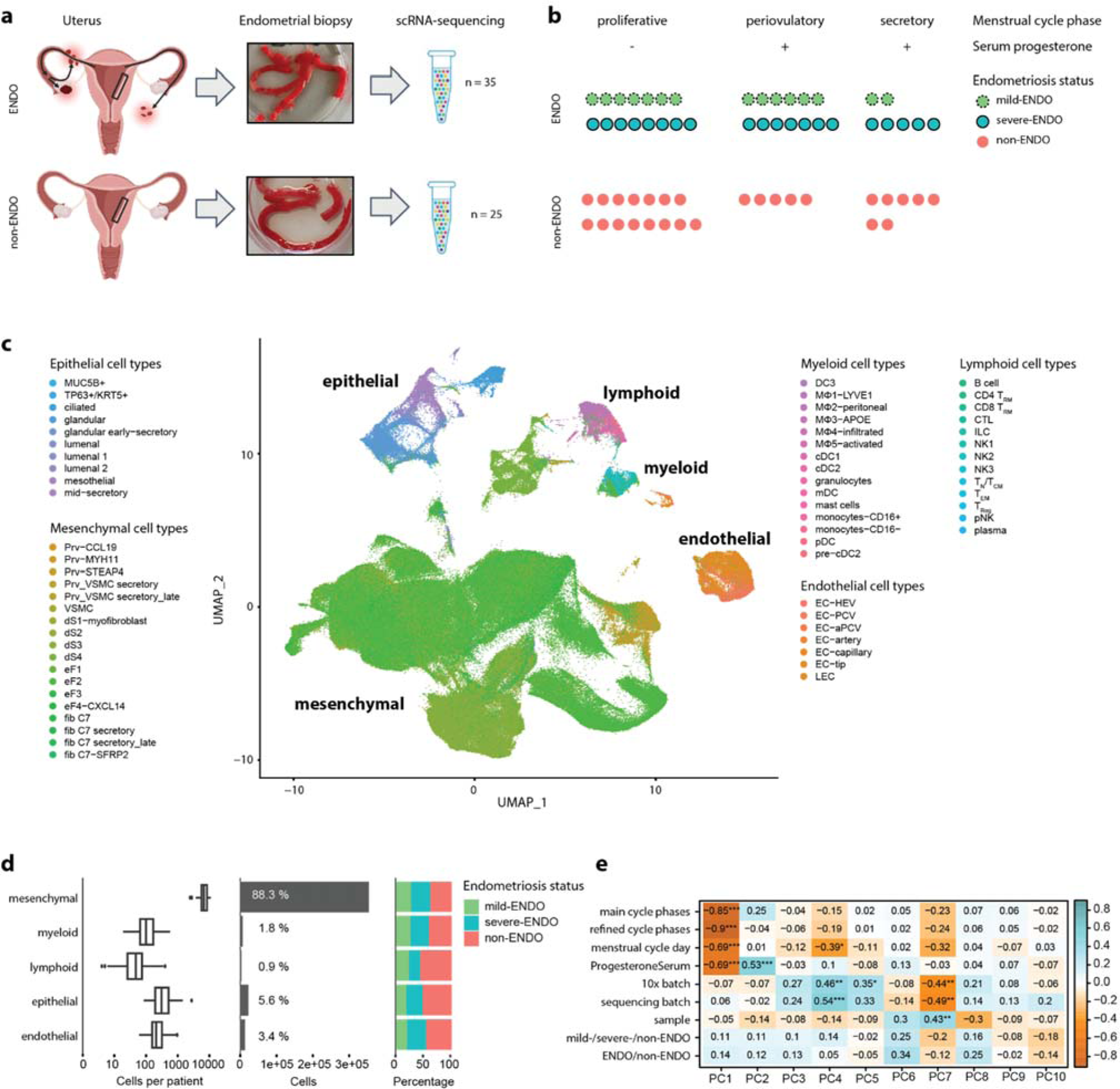
Single-cell atlas of the endometrium from women with and without endometriosis. **(a)** Schematic illustration of uterus (left), image of endometrial biopsies (middle), followed by single-cell RNA (scRNA)-capture with 10x Genomics and scRNA-sequencing with Illumina (right). The endometrial single-cell atlas includes 35 endometrial biopsies from women with endometriosis (ENDO) and 25 endometrial biopsies from women without endometriosis (non-ENDO). The black box in the uterus schematic delineates parts of the endometrium. The most widely accepted theory of endometriosis origin (Sampson, 1927) postulates the migration of shed endometrial tissue during menstruation to the peritoneal cavity (uterus schematic, black errors), where it forms endometriotic lesions (uterus schematic, red cell aggregates). **(b)** Distribution of samples in regard to menstrual cycle phase (proliferative, periovulatory and secretory) and endometriosis status (mild-ENDO, severe-ENDO and non-ENDO). Women with proliferative menstrual cycle phase did not have measurable serum progesterone (threshold = 2nmol/l). **(c)** UMAP of the entire endometrium single-cell atlas from women with and without endometriosis, colored by cell types (n=63). **(d)** Main cell type frequencies per patient (n=60, boxplot, left) and entire dataset (barplot, middle), with mesenchymal cells being the most abundant cell type. The overall cell number distribution between non-ENDO, mild-ENDO and severe-ENDO is balanced (barplot, right). **(e)** Eigencorplot showing the effect of different co-variants of the atlas dataset on principal components (PCs) with Pearson correlation values and multiple-testing adjustment with the Benjamini-Hochberg procedure. Menstrual cycle phase parameters significantly correlate with PC1, PC2 and PC4. In contrast, no significant correlation was identified between the endometriosis status and any of the initial ten principal components. Statistical significance symbols represent: *** <0.001, ** <0.01, * <0.05.

We successfully created a comprehensive single-cell atlas of the endometrium, comprising 283,789 ENDO and 182,582 non-ENDO patient cells. The samples had on average 7,773 cells, with 4,604 counts and 2,419 features per cell after SCT normalization, displaying no discernible difference between endometriosis status (Extended Data Fig. 1a,b). Using Symphony, we identified 57 cell types by alignment to a reference atlas specifically for endometrium and endometriosis^16,21^. The characteristic cell type marker genes were expressed as in the reference atlas^16^ (Extended Data Fig. 2). The cell type annotation was further refined to 63 cell types by graph-based clustering to accommodate changes during the menstrual cycle (Fig. 1c, Extended Data Fig. 1c,d, Methods). The cell types can be categorized into five main cell types: mesenchymal (88.3%), epithelial (5.6%), endothelial (3.4%), lymphoid (0.9%) and myeloid (1.8%) (Fig. 1d). The transcriptional variance within the dataset was captured through sample-wise Principal Component Analysis (PCA), revealing a significant correlation between the first Principal Components (PC1, PC2 and PC4) and key menstrual cycle phase parameters, including menstrual cycle phase, cycle day and serum progesterone levels (Fig. 1e). Surprisingly, no statistically significant correlation was identified between the endometriosis status and any of the initial ten Principal Components (Fig. 1e, Extended Data Fig. 1e,f and 3a). This suggests a minor impact of the endometriosis status on the endometrial transcriptional landscape, consistent with previous bulk-sequencing studies^11,12^. The nuanced influence of endometriosis on the molecular characteristics of endometrial tissue underscores the necessity of accounting for menstrual cycle phase, minimizing confounding factors, and exploring subsets of cells for a more comprehensive understanding of the relationship between endometriosis and the endometrial transcriptome. To mitigate confounding factors, we generated our single-cell atlas within the same laboratory, employing unvarying processing methods and encompassing detailed clinical data with menstrual cycle staging (Supplementary Table 1). This approach provides a powerful, unmatched resource for investigating endometrial biology and diagnostic signatures for endometrial pathologies like endometriosis.

### Cellular changes in the menstrual cycle drive transcriptional changes in the endometrium

Significant changes in cellular content, morphology, and transcription accompany the different menstrual cycle stages^22,23^. In the PCA plot of PC1/PC2 (Fig. 2a), proliferative/periovulatory and early-, mid-, and late-secretory samples cluster distinctly, underscoring the substantial impact of the menstrual cycle on the endometrial transcriptional landscape^23^, with PC1 and PC2 together accounting for 49.9% variance. Moreover, a clear visual separation in the UMAP between proliferative/periovulatory, early-, mid-, and late-secretory samples is evident, particularly in mesenchymal, epithelial, and endothelial cells (Fig. 2b). Interestingly, proliferative and periovulatory samples overlapped by PCA or UMAP (Fig. 2a,b), suggesting a similar transcriptional profile and a delayed effect of progesterone on endometrial transcription. Epithelial cells from early-, mid-, and late-secretory phases exhibited a clear gradient on the UMAP, prompting a pseudotime trajectory analysis with Monocle to infer the order of the samples within the menstrual cycle phases (Fig. 2c, Extended Data Fig. 3b).

**Fig. 2.**
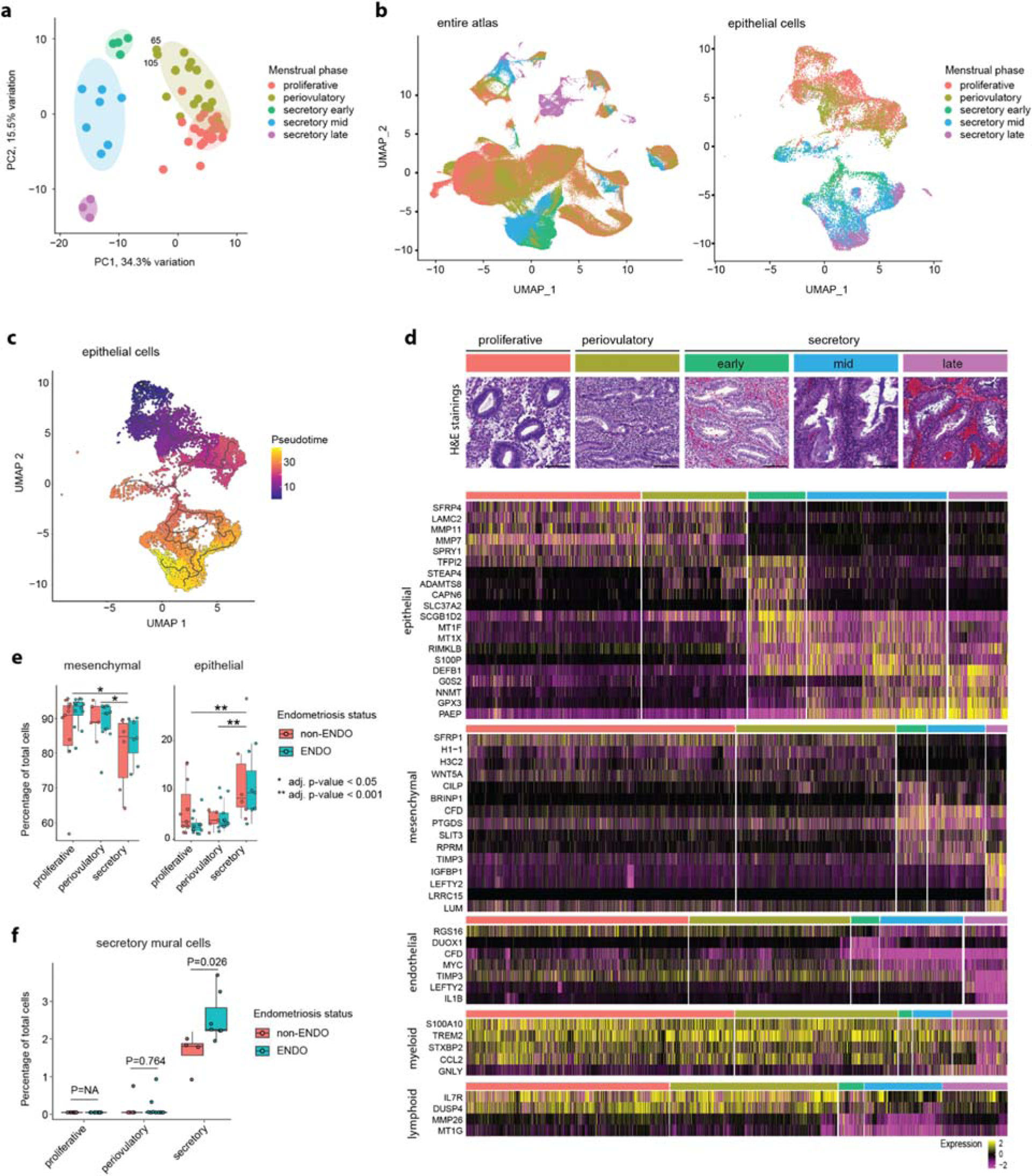
Cellular changes in the Menstrual Cycle drive transcriptional changes in the endometrium. **(a)** Scores plot of the first two principal components from PCA of sample-wise whole transcriptome. PC1 accounts for 34.4% of the total variance, while PC2 explains 15.5% (PCAtools). The proliferative and periovulatory samples do not cluster distinctly, whereas early-, mid- and late-secretory samples cluster separately, marked by ellipses (‘t’ distribution, 0.90 confidence interval, late-secretory ellipse added manually). Samples 65 and 105 (labeled) are in the transition between periovulatory and early secretory phases. **(b)** UMAPs of the entire dataset (left) and the re-integrated epithelial cells (right), colored by menstrual cycle phase, show a clear visual separation between proliferative/periovulatory, early-, mid-, and late-secretory samples, particularly in mesenchymal (endometrial fibroblasts, C7 fibroblasts, and mural cells), epithelial (glandular), and endothelial cells. **(c)** UMAP of the re-integrated epithelial cells, colored by pseudo time prediction from monocle3. **(d)** Top: Representative H&E staining from each key menstrual cycle phase (scale bar: 0.1mm). Bottom: Heatmap of selected marker gene expression with high log fold changes over the key menstrual cycle phases (proliferative, periovulatory, early-, mid-, and late-secretory). Samples were ordered by menstrual cycle phase and pseudo time. **(e)** Boxplots of mesenchymal and epithelial cell frequency as a percentage of total cells, split by menstrual cycle phase (stringent patient exclusion criteria, 32 ENDO, and 22 non-ENDO samples). P-values were calculated with a two-tailed Student’s t-test and were adjusted for multiple testing with the Benjamini-Hochberg procedure, * adj. p-value < 0.05, ** adj. p-value < 0.01. **(f)** Boxplot of secretory mural cell frequency as a percentage of total sample cells. Stringent patient exclusion criteria were applied and late secretory samples were excluded due to the imbalance in endometriosis status, resulting in 32 ENDO, and 20 non-ENDO samples. P-values were calculated with a two-tailed Student’s t-test and were adjusted for multiple testing with the Benjamini-Hochberg procedure.

To characterize the transcriptomic alterations occurring within the main cell types throughout the menstrual cycle, a comparative analysis with Seurat^24^ was conducted of the transcriptome across the key menstrual cycle phases. This investigation, stratified by the primary five cell types, uncovered 3643 genes to significantly change in expression over the menstrual cycle phases (adjusted p-value < 0.05, avg_log2FC > 2) (Supplementary Table 2), underscoring the substantial transcriptomic variations during the menstrual cycle (Fig. 1d). The observed expression changes of menstrual marker genes mirrored the sequence outlined in a comprehensive C1 menstrual cycle atlas from healthy volunteers with a regular cycle^23^. Notably, retrospective aligning of our atlas samples to their menstrual cycle phases^23^, using Symphony^21^, resulted in perfect correspondence (Extended Data Fig. 3c-f). This alignment instills a high level of confidence in the accuracy of our menstrual cycle staging. Our single-cell atlas, with almost 50 times more cells per patient and detailed annotation, establishes a novel reference point for diverse datasets and offers unprecedented insights into the intricacies of menstrual cycle dynamics. Investigation into cell type frequency changes between menstrual cycle phases reveals significant shifts. Generally, epithelial cells increase, while mesenchymal cells decrease from proliferative to secretory phase (Fig. 2e, Extended Data Fig. 4a). Approximately a third of the refined cell types show significant abundance changes during the menstrual cycle (Supplementary Table 3). Examining cell type frequency differences between ENDO and non-ENDO, the only significant abundance change was observed in secretory mural cells (Prv_VSMC secretory) (unadjusted p-value = 0.009, Extended Data Fig. 4b). Mural cells are important for vascular development and stability. The significant 1.6-fold increase of secretory mural cells in ENDO in comparison to non-ENDO (adjusted p-value = 0.026, Fig. 2f,) suggests increased vascularization in the secretory endometrium of endometriosis patients. In line, an increase of perivascular endometrial mesenchymal stem cells has been detected in menstrual fluid^25^, supporting the ‘stem cell hypothesis’ of endometriosis pathogenesis^26^.

### Transcriptional alterations in the endometrium of women with endometriosis

Next, we investigated whether transcriptional changes can be observed in the endometrium of women with endometriosis, and if these changes can be associated with processes suggested to contribute to endometriosis susceptibility^9^. To detect robust alterations in transcription between endometriosis status, we focused on the homogeneous proliferative menstrual cycle phase (Fig. 2)^22,23^.

Cluster-wise differential gene expression analysis revealed 427 unique differentially expressed genes (DEGs, p-value < 0.05) between non-ENDO and mild-, severe- or all-ENDO using Muscat^27^ (Fig. 3a, Supplementary Table 4, Extended Data Fig. 5a). Notably, severe-ENDO exhibited a fivefold increase in DEGs compared to mild-ENDO (Fig. 3a). Cluster- and stage-wise functional enrichment analysis identified significant perturbations in inflammation, cellular adhesion, cell proliferation and survival, angiogenesis, and the nervous system, as detailed in the following sections (Fig. 3b, Extended Data Fig. 5b, Supplementary Table 5).

**Fig. 3.**
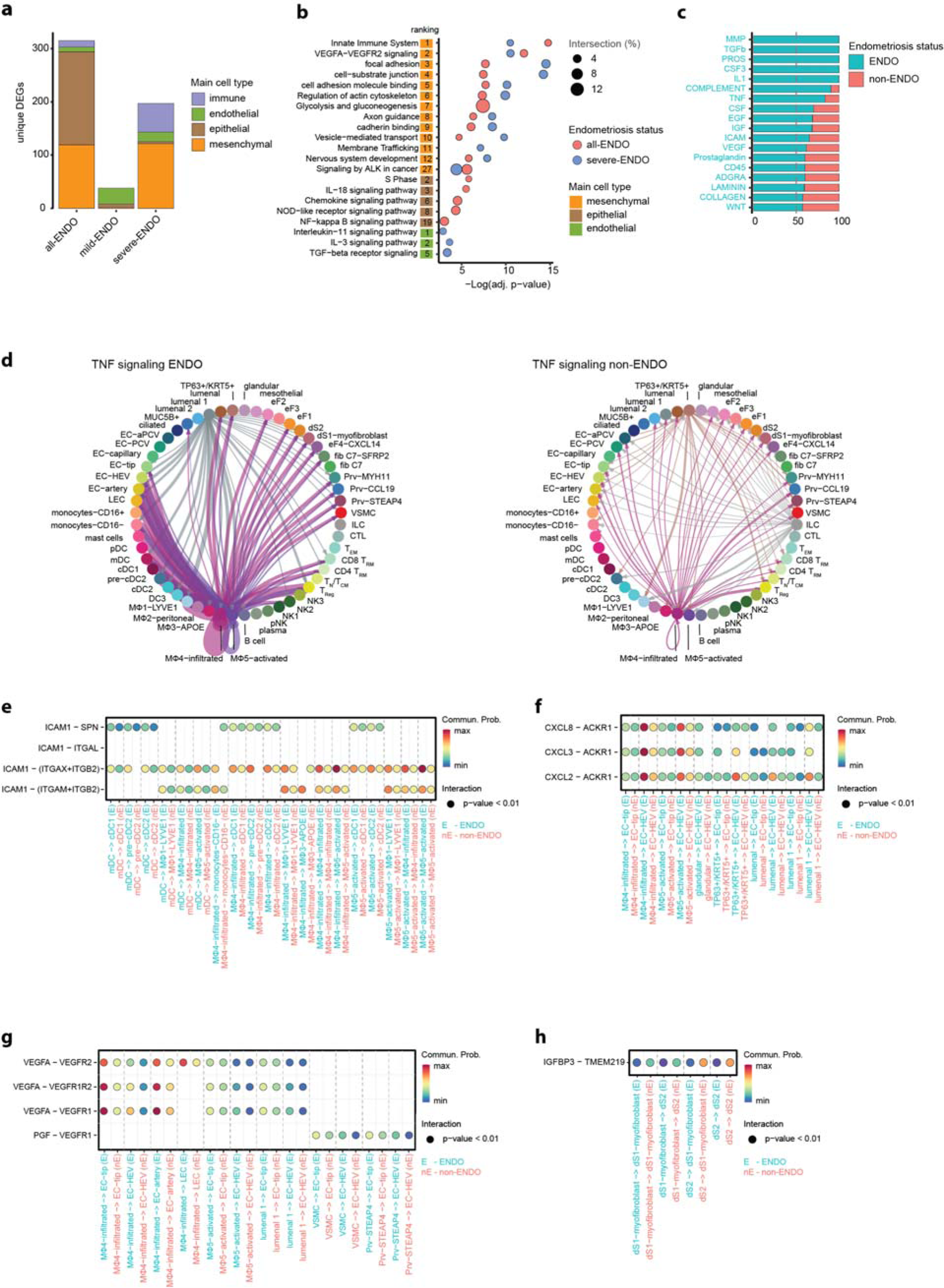
Inflammation, Adhesion, Proliferation, and Angiogenesis pathways and ligand-receptor interactions are upregulated in endometriosis endometrium of the proliferative phase. **(a)** Barplot showing number of unique differentially expressed genes (DEGs, p-value < 0.05, average expression > 18.53202) between non-ENDO (n = 11) and all-ENDO (n = 12), mild-ENDO (n = 4) and severe-ENDO (n = 8) samples within immune, endothelial, epithelial, and mesenchymal cell types, analyzed by Muscat. DEG analysis identified 427 unique DEGs between non-ENDO and mild-, severe- or all-ENDO (38, 192, and 309 DEGs, respectively). Compared to non-ENDO, there were 5 times more in severe-ENDO than in mild-ENDO. The lymphoid and myeloid immune cells of severe-ENDO displayed 54 DEGs, in stark contrast to the absence of any immune DEGs in mild-ENDO. **(b)** Selected terms enriched in the cluster- and stage-wise functional enrichment analysis of DEGs, with term sizes between 25 and 750. The rank by adjusted p-value noted under ranking. **(c)** Selected signaling pathways with overall enriched activity in ENDO (n = 12) in comparison to non-ENDO (n = 11). **(d)** Circle plot of TNF signaling pathway between all cell types in ENDO (left, n = 12) and non-ENDO (right, n = 11). **(e-h)** Bubble plots of selected communications with significant interaction probabilities (unadjusted p-value < 0.01) within the ICAM (e), CXCL (f), VEGF (g) and IGFBP (h) signaling pathways for ENDO (n = 12) and non-ENDO (n = 11). Further p-value adjustment is not recommended by the CellChat author.

Using CellChat^53^, we quantitatively analyzed intercellular communication networks, revealing overall enriched activity of several ligand-receptor interactions in the endometrium of women with ENDO, including TGFb, IL1, TNF, ICAM, VEGF, and WNT. These interactions suggest stimulated inflammation, angiogenesis, proliferation, and cell survival as discussed in more detail below (Fig. 3c, Extended Data Fig. 5c,d).

These findings show that regulatory disturbances are already present in the endometrium of women with endometriosis and are more pronounced in severe endometriosis cases. Functional enrichment and ligand-receptor analysis highlight dysregulation of inflammation, cell proliferation, survival, adhesion, angiogenesis, axonal guidance and WNT/NOTCH pathways, consistent with the processes thought to be required for endometrial cells to establish viable lesions at ectopic sites. Enhanced interactions in ENDO between macrophages and other cell types point toward a pivotal role for macrophages. These findings provide novel insights into the molecular intricacies of endometriosis, which occur early in development and offer potential targets for therapeutic intervention.

### Inflammation and adhesion are enhanced in ENDO endometrium

Macrophages play a central role in normal endometrial tissue remodeling and endometriosis lesions, contributing to growth, development, vascularization, and lesion innervation^54^. Endometrial macrophages were proposed to favor endometriosis lesion formation, however, the mechanism is convoluted^55^. We observed a significantly enhanced interaction between macrophages and stromal, endothelial, epithelial and immune cells through TNF – TNFR1/2 signaling (Fig. 3d, Extended Data Fig. 5e). Binding of tumor necrosis factor (TNF) to TNF receptor 1 (TNFR1) activates NFκB and mitogen-activated protein kinase (MAPK) signaling pathways, responsible for regulating transcription of pro-inflammatory mediators including cytokines, chemokines and the atypical chemokine receptor ACKR1^31,56,57^. TNF is also deregulated in endometriosis lesions^29^. Furthermore, there is a strongly enhanced interaction in ENDO between macrophages, dendritic cells, and epithelial cells with multiple immune cells mediated through ICAM1/2 and its receptors (Fig. 3e). ICAM expression is strongly triggered in epithelial and immune cells under inflammatory stimulation and TNF-inducible ICAM1 regulates the recruitment of circulating leukocytes to sites of inflammation^58,59^. Moreover, we observed an enhanced interaction between macrophages and epithelial cells with endothelial cells through the chemokines CXCL2, CXCL3, and CXCL8 with ACKR1 (Fig. 3f), known to regulate inflammatory response.

The complex interplay among different cell types through secreted cytokines is critical in the inflammatory microenvironment^28^. Our analyses further revealed increased cytokine expression (CXCL2, CXCL3, TGFB1, CXCL12) and upregulated IL-3, IL-11 and IL-18 signaling pathways in ENDO (Fig. 3b, Supplementary Table 5). NFκB pathway and JAK1, previously implicated in endometriosis lesions^29,30^, were both significantly higher expressed in ENDO endometrium (Fig. 3b, Supplementary Table 5). NFκB orchestrates inflammatory responses by promoting the synthesis of pro-inflammatory cytokines and chemokines, such as IL1, IL6, IL8, TNFa, RANTES, MIF, and ICAM1, while JAK1 regulates the immune system and proinflammatory response^31,32^. Moreover, the upregulation of CXCL12 in mesenchymal cells, which in bone marrow supports the maintenance of hematopoietic stem cell niche^33^ and may play a role in the stem/progenitor nature of endometrial mesenchymal cells^34^, could increase immature endometrial cells and thereby endometriosis susceptibility.

Upon entry into the peritoneal cavity, adhesion of refluxed endometrial cells facilitates attachment to the underlying peritoneal tissue and contributes to migration and invasion^9^. Several genes involved in focal adhesion and cytoskeleton reorganization, including ACTB, RHOA, COL4A2, COL5A2, JAK1, and MYH9, were upregulated in mesenchymal cells (Fig. 3b, Extended Data Fig. 6a), in line with stomal cells leading invasion^35^. Further upregulated in ENDO mesenchymal cells was TGFB1|1, shown to promote focal adhesion, migration, and EMT in carcinoma^36^.

### Proliferation and cell survival are enhanced in ENDO endometrium

We observed increased expression of genes associated with cell proliferation and survival pathways, including NFκB signaling and ALK signaling in ENDO epithelial and mesenchymal cells (Fig. 3b) as well as decreased cell death receptor signaling^31,37,38^. NFκB, known for its role in cell proliferation, apoptosis suppression, and adhesion molecule expression, is overactive in endometriotic lesions^29^. Its activation by NOD-like receptors, also significantly enriched, underscores its importance^31^. NFκB inhibitors show potential as non-hormonal drugs against endometriosis, reducing inflammation, angiogenic factors, and MMPs in vitro and lesion size in animal models^29,39^. The ALK signaling pathway, among the top 30 enriched terms in mesenchymal cells, mediates signaling through various pathways, leading to increased cell proliferation, cell survival, and cell cycle progression^40^. Ligand-receptor analysis further indicates reduced apoptosis in ENDO (Fig. 3h) by decreased signaling between the cell death receptor TMEM219 and IGFBP3^63^ in several mesenchymal cell-cell interactions in ENDO (Fig. 3h). Enhanced proliferation, enhanced cell survival and decreased apoptosis may contribute to effective endometriosis lesion formation^31,37,38^.

### Angiogenesis is enhanced in ENDO endometrium

Angiogenesis is essential for sustaining endometriosis lesions by ensuring nutrient supply^9^. VEGFA-VEGFR2 signaling (Fig. 3b) and several angiogenesis-mediating ligand-receptor interactions were enhanced in the ENDO (Fig. 3g). The interaction between macrophages and endothelial cell types is increased through VEGFA-VEGFR1/-VEGFR1R2/-VEGFR2 in ENDO. VEGFA-VEGFR2 is the most prominent ligand-receptor complex in the VEGF system^60^ and genetic variants in the VEGFR2 gene (KDR) have previously been associated with endometriosis susceptibility^61^. Furthermore, an enhanced interaction in ENDO between mural cells (perivascular (Prv-STEAP4) and vascular cells (VSMC)) and endothelial cells (EC-tip, EC-HEV) through the angiogenesis promoting PGF-VEGFR1 ligand-receptor binding^62^ was observed (Fig. 3g). VEGF signaling through VEGFR1/VEGFR2 ultimately promotes endothelial cell proliferation, migration, survival, and vascular permeability^60^. Compellingly, the abundance of secretory mural cells is significantly higher in ENDO than in non-ENDO (Fig. 2f), which aligns with an increase in endothelial cell proliferation through enhanced VEGF signaling.

Our analysis further unveiled significant enrichment of TGFb signaling in endothelial cells, by upregulation of genes such as TGFB1 and JAK1 (Extended Data Fig. 5b). TGFb is an inflammatory growth factor that regulates cell adhesion, invasion, and angiogenesis^41^, and was found upregulated in endometriosis lesions^9^. In tumors, TGFb induces the metabolic conversion of glucose to lactate via glycolysis, a process referred to as the “Warburg effect”, with lactate increasing angiogenesis and cell invasion^42^. Glycolysis and gluconeogenesis was a significantly enriched term in ENDO mesenchymal cells (top 7; Fig. 3b) suggesting a possible occurrence of a Warburg-like effect, in line with a peritoneal endometriosis report^43^. Dysregulation of TGFb signaling pathway contributes to fibrosis and neovascularization in various diseases, underscoring its therapeutic potential in endometriosis management^44,45^.

In summary, we observed enhanced angiogenesis signaling and ligand-receptor interactions as well as an increase of secretory mural cells in the endometrium of women with endometriosis, which may explain increased endometriosis susceptibility.

### Nervous system and WNT/NOTCH in ENDO endometrium

Painful symptoms associated with endometriosis lesions are proposed to arise from interactions with nerve fibers at ectopic sites^41,42^, mediated by both immune and mesenchymal-derived inflammation^43,44^ which stimulate sensory nerve fiber infiltration into established lesions^45^. However, controversy exists regarding the increase in sensory nerve fibers in the endometrium of women with endometriosis^43^. Axon guidance and nervous system development were among the top 20 enriched terms (Fig. 3b) in ENDO mesenchymal cells. Among the differentially expressed genes was the chemokine CXCL12 (SDF-1), which plays a critical role in axon pathfinding and elongation potentially via the SDF-1α/Rho/mDia pathway^51^. This suggests a possible augmentation of nerve fibers in the endometrium of women with endometriosis.

The endometrium undergoes monthly regeneration driven by mesenchymal stem and epithelial progenitor cells and relies on programmed differentiation and maturation for normal function. We detected a subtle increase in overall WNT signaling (WNT2/3/7A - (FZD3/4/6+LRP6)) and a marginal increase in overall NOTCH signaling in ENDO (Fig. 3c, Extended Data Fig. 5c). Notably, cross-talk between NOTCH and WNT pathways affects epithelial cell differentiation, with increased WNT activity leading to enhanced ciliary commitment^64^, a feature observed in endometriosis lesions^38^.

### Accurate prediction of endometriosis based on all cells or cycling fibroblasts using ScaiVision

Next, we trained diagnostic models for endometriosis based on gene expression changes within the endometrium. We employed the machine learning algorithm CellCnn via Scailyte’s ScaiVision platform, which can assess and integrate expression changes of hundreds of genes simultaneously and employ representation learning to discover relevant biomarkers for the disease model^65^.

To ensure the reliability of predictions, ScaiVision models were trained exclusively on proliferative phase samples (Supplementary Table 1) (Fig. 2d). Employing a 5-fold Monte Carlo cross-validation scheme^66^, samples were allocated for training (56%), validation (24%), and evaluation (20%), resulting in 25 training and validation datasets, and 5 independent evaluation datasets (Fig. 4a, Extended Data Fig. 6a-c). We used four different feature selection methods to reduce the dimensionality of the data prior to model training: 1) PCA identifying components of the highest variance^67^, 2) CorrgPCA using correlation-based PCA (Methods), 3) diffgen based on non-parametric Wilcoxon rank sum test^68^ and 4) hvg identifying highly variable genes^24^. All methods exhibited a median validation AUC of 0.78, with PCA, CorrgPCA, and diffgen achieving median validation AUC of ≥ 0.78 in 3 of the 5 folds (Fig. 4b, top). The top three validation models from each fold performed well on the independent evaluation datasets, resulting again in median evaluation AUCs ≥ 0.78 in 3 of 5 folds for PCA and 2 of 5 folds for corrgPCA (Fig. 4b, bottom). Our presented models demonstrate equal efficacy in predicting both mild and severe endometriosis, highlighting prediction robustness for different endometriosis severity (Extended Data Fig. 6d).

**Fig. 4.**
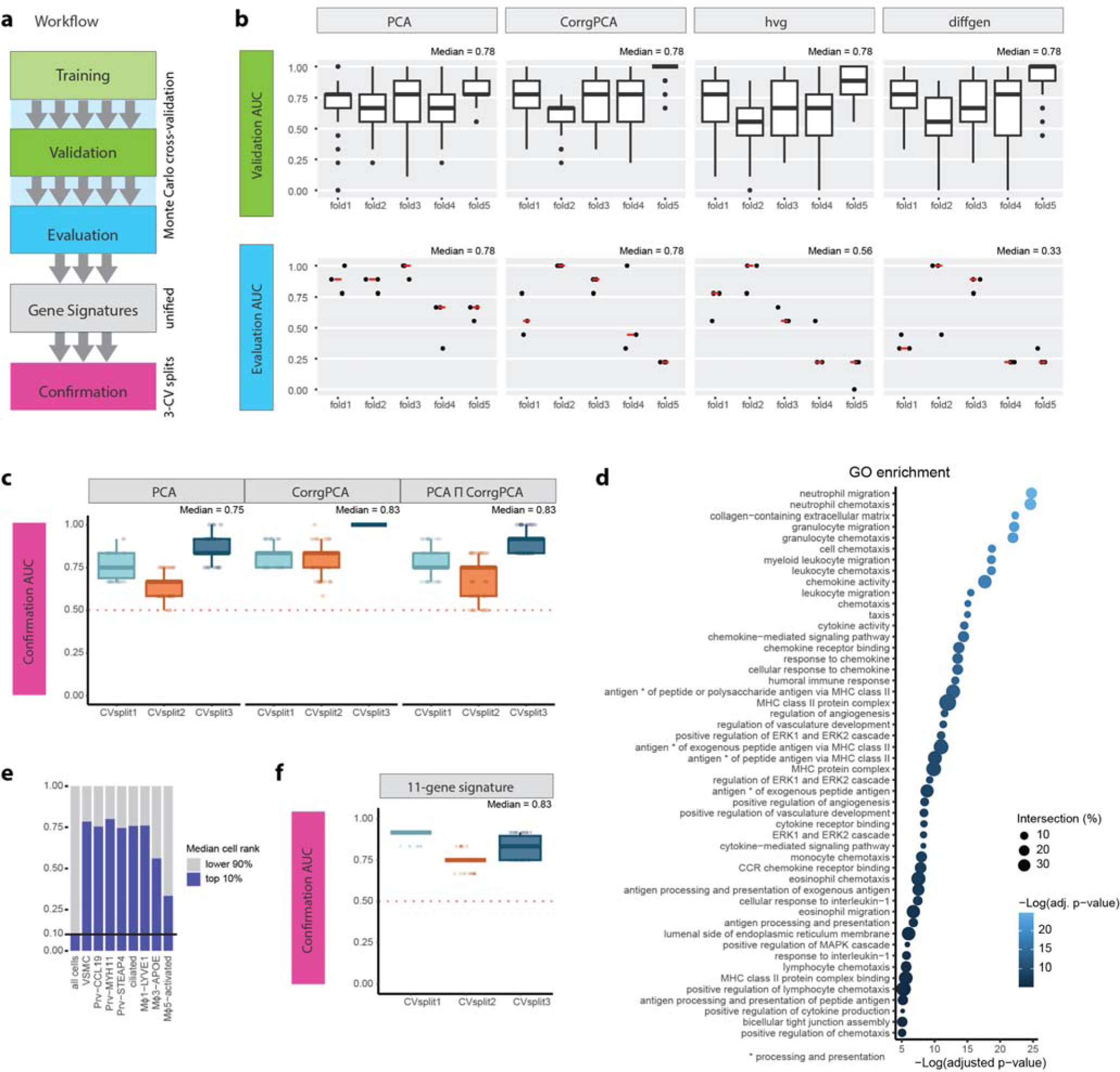
Prediction of endometriosis with ScaiVision using convolutional neural networks. **(a)** Workflow outlining the ScaiVision model training and gene signature confirmation process. Proliferative samples (n = 25) were allocated for training and validation (76%), and evaluation (24%) using a 5-fold Monte Carlo cross-validation scheme. The training and validation samples were further stratified with 5-CV splits, resulting in 25 training and validation datasets. Unified gene signatures were derived by selecting genes with the top 20% deepLIFT and integrated gradients importance scores and evaluated on 3-CV splits of all proliferative samples. **(b)** Boxplots illustrate the validation AUCs (n = 250 per fold and feature selection method) and evaluation AUCs (n = 3 per fold and feature selection method), categorized by feature selection methods (PCA, CorrgPCA, hvg, and diffgen). The box centerline represents the median AUC for validation data, while the dark red line signifies the median AUC for evaluation data. The median AUC per plot is displayed on its top right. PCA and CorrgPCA demonstrate superior performance in evaluation, achieving median AUCs of 0.78. **(c)** Boxplots of the confirmation AUC (n = 50 per box plot) for the unified PCA and CorrgPCA gene signatures and their intersect (PCA ∩ CorrgPCA), showcasing median AUCs consistently above 0.7 for each CV split. **(d)** Top 50 GO terms and pathways, ranked by the adjusted p-value, highlight differences in the immune system, extracellular matrix, angiogenesis and the ERK1/2 and MAPK cascade in the endometrium of women with endometriosis. Term sizes below 10 and above 500 are excluded. **(e)** Barplot showcases cell types with more than 25% of cells within the top 10% of filter scores for predicting no endometriosis. Specific subsets of mural cells (VSMC, PRE-CCL19, Prv-MYH11, Prv-STEAP4), ciliated epithelial cells, and myeloid cells (Mϕ1, Mϕ3, and Mϕ5) exhibit increased filter response scores. **(f)** Boxplot of the confirmation AUC (n = 150 per box plot) for the 11-gene signature, derived from the intersection between the ScaiVision-derived endometriosis signatures (PCA and CorrgPCA) and the top 300 cluster-wise DEGs.

**Fig. 5.**
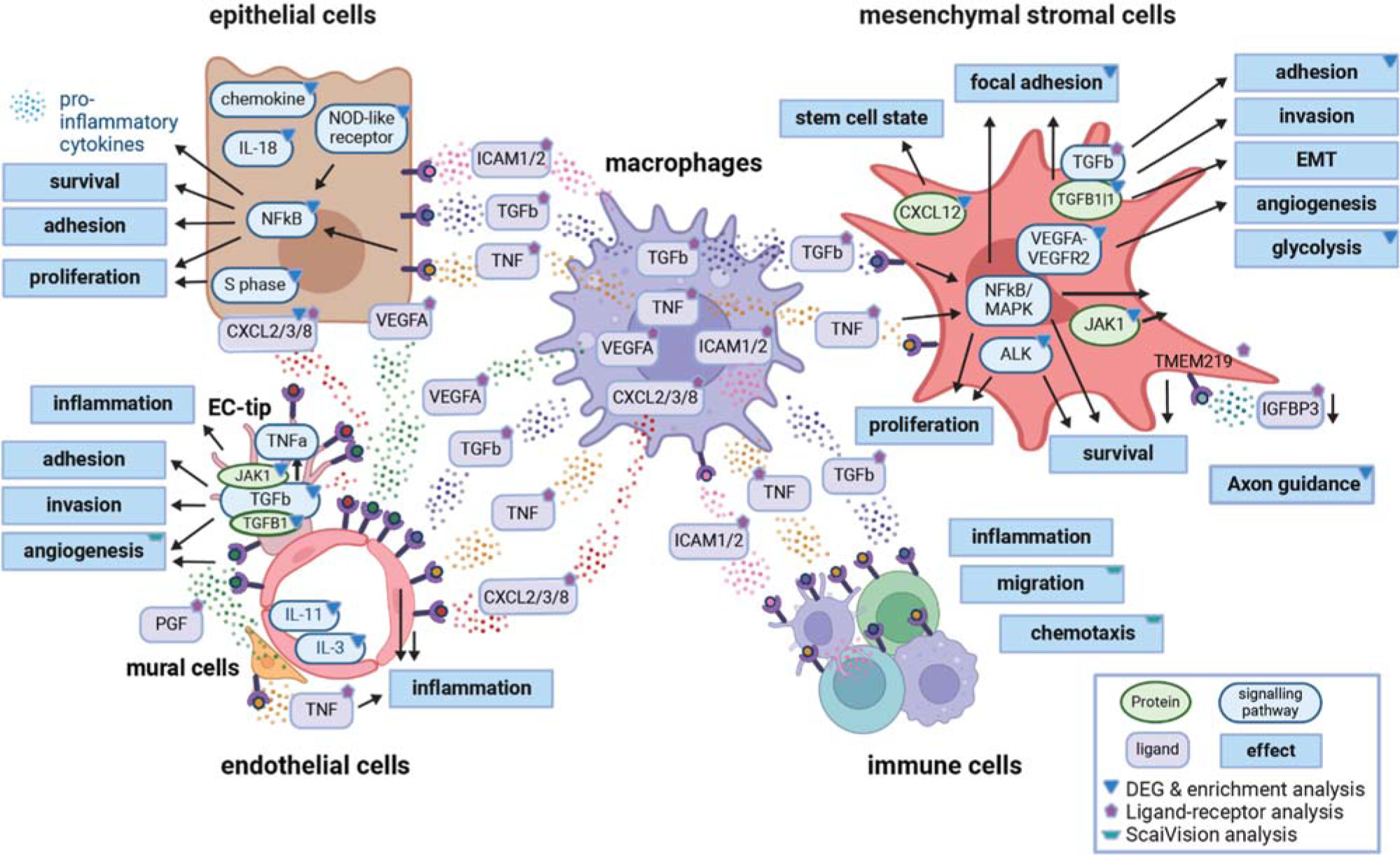
Graphical summary of enhanced ligand-receptor interactions and pathways in the endometrium of women with endometriosis. Ligand-receptor analyses point toward a pivotal role for macrophages in endometriosis endometrium. Enhanced interactions between macrophages and various endometrial cell types, mediated by TNF, CXCL, TGFb, ICAM1, VEGF and IGFBP signaling, indicate elevated inflammation, angiogenesis, proliferation, and cell survival. Additionally, elevated signaling between epithelial and endothelial cells via CXCL and VEGF pathways was observed. Functional enrichment analysis of differentially expressed genes (n > 400), stratified by cell-types, underscores significant perturbations in inflammation (chemokine signaling, NOD-like receptor signaling, NFκB signaling), cellular adhesion (TGFb receptor signaling, adhesion, extracellular compartment), cell proliferation and survival (NFκB signaling, ALK signaling), angiogenesis (TGFb and VEGFA-VEGFR2 signaling), and nervous system processes (development and axon guidance) across multiple cell types. Furthermore, ScaiVision analysis underscores significant changes in cytokine and chemokine activity, chemotaxis and migration of various immune cells and angiogenesis. These findings are consistent with the processes believed to be crucial for endometrial cells to establish viable ectopic lesions. In conclusion, the generation and analysis of our single-cell atlas reveals at unprecedented cell-type resolution that regulatory disturbances are present already in the endometrium of women with endometriosis, potentially increasing endometriosis susceptibility and suggesting therapeutic targets for intervention.

Next, we extracted the genes responsible for driving the performance of the top networks in the 5-fold from the well-performing PCA and corrgPCA feature selection methods based on deepLIFT and integrated gradients importance scores (see Methods, cite deepLIFT and captum). The unified PCA (215 genes) and CorrgPCA (210 genes) signatures shared over 60% of the genes (PCA∩CorrgPCA), validating the selection process of predictive genes (Extended Data Fig. 6e, Supplementary Table 6). Cross-validation of the unified PCA, CorrgPCA and PCA∩CorrgPCA gene signatures with 3 Cross-Validation (CV) splits of the proliferative samples consistently achieved high AUCs, with the best performing networks of each split achieving an AUC of 0.75, 0.83 and 0.83 respectively (Fig. 4c). This result confirms the importance of the ScaiVision-selected gene signatures for the prediction of endometriosis.

Subsequently, we aimed at predicting endometriosis based on only a subfraction of cells, which could be experimentally isolated from the endometrial tissue. In the proliferative phase samples, endometrial fibroblasts with high G2M values (cycling fibroblasts) account for 11.6% of the endometrium (Extended Data Fig. 7a). There are 32% more cycling fibroblasts in ENDO (13.0%) than non-ENDO (9.8%) (non-significant), in line with a previous study^16^. The ScaiVision models trained on cycling fibroblasts could predict endometriosis with high accuracy, achieving an evaluation AUC of ≥ 0.78 on at least 3 of the 5 folds for diffgen, corrgPCA, and PCA (Extended Data Fig. 7b,c). The united PCA, CorrgPCA, and PCA∩CorrgPCA signatures achieved high accuracies and median AUCs of 1.0, for the best-performing networks of each 3 Cross-Validation (CV) splits (Extended Data Fig. 7d). GO term analysis revealed an enrichment of structural terms (collagen and ECM), apoptosis, myeloid cell differentiation, growth factor binding, MHC and responses to TNF (Extended Data Fig. 7e). These data emphasize the predictive potential of ScaiVision for endometriosis on only a fraction of endometrial cells, additionally highlighting the importance of cycling fibroblasts in endometriosis.

In conclusion, we successfully trained diagnostic models for endometriosis based on all cells or on cycling fibroblasts only and extracted unified biomarker signatures from the PCA and CorrgPCA models.

### ScaiVision predicts endometriosis based on multiple cell types and genes in immune pathways and angiogenesis

The unified PCA gene signature from all cells (Supplementary Table 6), analyzed via Gene Ontology (GO) terms, highlights altered cytokine and chemokine activity, response, binding and signaling pathway. Additionally, chemotaxis and migration of various immune cell types were enriched, underscoring endometriosis-related immune system changes in endometrial tissue, in line with the DEG and receptor-ligand analysis (Fig. 3b,f). Regulation of angiogenesis and vasculature development were among the top 25 enriched GO terms, in line with the DEG and receptor-ligand analysis (Fig. 3b,g). Furthermore, collagen-containing extracellular matrix is top-ranked (rank 3, Fig. 4d), in line with the DEG pathway enrichment analysis (Supplementary Table 5). The key genes in the unified PCA signature by deepLIFT and integrated gradients importance scores, namely IL1B, CCL4, CCL3, HLA-DRA, HLA-DPA1, CD74, A2M, and ACKR1, play crucial roles in immune responses and DUSP2 dephosphorylates MAPKs (Supplementary Table 6).

Utilizing CellCnn’s strength in identifying cell subsets associated with the disease, ScaiVision identified mural cells, macrophages, and ciliated epithelial cells to be crucial for non-ENDO prediction (Fig. 4e, Supplementary Table 7) and mural cells, dendritic cells, and several epithelial cell types for ENDO prediction (Supplementary Table 7). This is in line with the many altered gene expression changes and ligand-receptor interactions between myeloid, epithelial, and endothelial cells (Fig. 3), as well as increased VEGF signaling in mural cells with subsequent increased cell frequency (Fig. 3, Extended Data Fig. 4).

Reducing the ScaiVision-derived PCA and corrgPCA signatures, derived from all cells, through the intersection with the top 300 cluster-wise DEGs yielded an 11-gene signature (CCN1, CXCL2, CXCL3, SLPI, DIO2, MGP, COL4A2, ADAMTS1, CALD1, CEBPD, TPM1). ScaiVision models based on this 11-gene signature achieved high AUC for endometriosis prediction (median AUC = 0.83, Fig. 4f). Each gene in this signature plays a distinct role in biological processes. CCN1 promotes adhesion, migration, proliferation, and angiogenesis. Transcription factor CEBPD acts in immune responses. The chemokines CXCL2 and CXCL3, showing increased interaction with ACKR1 in ENDO (Fig. 3f), contribute to immunoregulation. SLPI protects against serine proteases, COL4A2 is linked to tissue remodeling, and DIO2 is crucial for thyroid hormone activity with previous endometriosis association^69^. ADAMTS1 has anti-angiogenic activity, is associated with inflammation, and has a metalloproteinase domain. MGP, CALD1, and TPM1 are associated with inhibiting ectopic tissue calcification, regulating smooth muscle and non-muscle contraction, and controlling muscle contraction, respectively. This refined gene signature provides insights into cellular processes, suggesting potential targets for further investigation.

## Discussion

The endometrium is a fundamental component of the female reproductive system for embryo implantation and growth. Anomalies in endometrial function are associated with multiple reproductive disorders, including endometriosis, a prevalent and debilitating disease^1,7^. The eutopic endometrium is thought to be the tissue of origin for endometriosis^8^. The underlying factors contributing to the onset of endometriosis and whether observable changes occur in the eutopic endometrium of affected patients are still controversial^10^. Our study addresses previous limitations by presenting the largest endometrium single-cell atlas from endometriosis (n = 35) and non-endometriosis (n = 25) patients (1) without hormonal treatment, (2) from different menstrual cycle phases, (3) analysis stratified by the donor’s menstrual cycle phase, (4) whole single-cell sequencing, (5) processed uniformly, and (6) accompanied by well-defined clinical data. By addressing these limitations, we identified significant gene expression changes and altered receptor-ligand interactions in the endometrium of endometriosis patients, potentially enhancing lesion formation and deepening our understanding of endometriosis pathophysiology. Applying this data to powerful neural network models we can accurately predict endometriosis based on endometrial biopsies.

The endometrium poses specific challenges in its study due to its monthly renewal cycle accompanied by expression changes of hundreds of genes^23^. Endometrial maturation is initiated by endometrial epithelial progenitor and mesenchymal stem cells in the basal layer, which proliferate under estrogen to repopulate the functional layer^70^. The rise of progesterone during the periovulatory and secretory phases prompts significant functional and morphological changes, as well as lineage specialization of the epithelial cells and decidualization of the stromal cells^71^. Dynamic immune cell infiltration supports this process, aiding tissue function transition and angiogenesis^54^. Our atlas, with refined menstrual cycle evaluation, detailed cell annotation, and high cell count per donor, maps cellular content and transcriptional changes across the menstrual cycle. It will serve as a crucial resource for an improved understanding of endometrial development, describing changes in cellular content, maturation, and intercellular relationships throughout the stages of the endometrial cycle. This insight is pivotal for comprehending how abnormalities in endometrial maturation may contribute to endometrial pathologies.

During the menstrual cycle key mechanisms and pathways undergo dynamic changes, including angiogenesis/VEGF^72^, immune functionality^23^, wound healing^54^, adhesion^73^, WNT and NOTCH^64^ and TGFb signalling^64,74^. External hormonal treatment can have diverse effects on menstrual cycle phase markers, rather than synchronizing them (Extended Data Fig. 3g,h). Therefore, it is essential to stratify endometrial analysis by the donor’s menstrual cycle phase and ensure an adequate number of donors for accurate interpretation of disease-specific changes, considerations often overlooked in previous studies.

Single-cell studies on endometriosis provided valuable insights ^14–19,34,38^. However, when comparing eutopic endometrium between endometriosis and non-endometriosis patients, these studies were limited by small eutopic sample sizes (n =< 3)^16^, inclusion of hormonally treated donors^16,17^, restricted cell subset analyses^19,34,38^ and/or neglect of stratification by the donor’s menstrual cycle phase^34^. In contrast, our atlas overcomes these issues with a large sample size (n = 60), exclusion of hormonally treated donors, comprehensive single-cell annotation and analysis, and studious stratification by the donor’s menstrual cycle phase.

This strategy enabled us to discover several differentially expressed genes in many cell types between non-ENDO and ENDO patients within the morphologic and transcriptionally homogeneous proliferative phase of the menstrual cycle^22,23^. The added refinement accompanying the examination of gene expression at the single-cell level confirms the advantages of a single-cell sequencing approach to studying this complex tissue. Our findings indicate that the gene expression changes particularly in inflammation, cellular adhesion, proliferation, survival, and angiogenesis pathways that occur in individual and complementary cell types may contribute to endometriosis susceptibility, offering hope for improved understanding and more effective treatments. In conclusion, this study reveals that endometriosis-related differences in gene transcription can be detected early in endometrial development, within the proliferative endometrium, paving the way for both targeted treatments and preventative measures against endometriosis.

Gene expression changes provided the foundation for developing in vitro diagnostic models based on ScaiVision, showcasing the potential of leveraging advanced computational techniques in unraveling complex molecular signatures. Endometriosis affects approximately 190 million women globally, causing significant disability and negatively impacting quality of life. An in vitro diagnostic could reduce the economic burden on healthcare systems, estimated to be over $10,910.96 per woman per year^2^, and reduce the delayed diagnoses leading to increased costs and preventing patients from adequate treatment. Particularly mild endometriosis is hard to detect through clinical examination or biomarkers^5^. Our single-cell-based models can predict both mild and severe endometriosis with accuracy, for hypothesis-generation without external validation and the discovery of single-cell signatures with predictive powers for endometriosis. While powerful single-cell sequencing is still expensive, our ongoing efforts involve translating our best-performing single-cell signature to a more cost-effective qPCR approach. In conclusion, our research offers a significant step forward in understanding endometriosis at the single-cell level, providing a foundation for improved diagnostics and targeted therapies.

## Methods

### Study approval and patient cohort

Patients were recruited at the Frauenklinik, Bern after the approval of this study by the Swiss Ethic Committee (KEK-BE 01780, 2019). Inclusion criteria for this study include women who provide Informed Consent, were above 18 years old and scheduled for laparoscopic surgery for reasons including symptoms of endometriosis, tubal ligation, salpingectomy, hysterectomy, idiopathic infertility, pelvic pain or other gynecological pathologies as part of their planned clinical treatment. Patients with pre-existing inflammatory diseases, malignancy, pregnancy, lactating or the onset of the last period more than 35 days before the surgery were excluded. Patients without endometriosis but with adenomyosis were excluded. Patients filled out a pain questionnaire before laparoscopy. Immediately before surgery, an endometrial biopsy was obtained using Pipelle-de-Cornier (Laboratoire CCD, 1103000) and peripheral blood was taken with an S-Monovette tube (Sarstedt, 04.1926.001). During laparoscopy, each patients’ pelvic cavity was visually inspected for endometriotic lesions. Endometriosis was confirmed for each endometriosis patient in this study by histopathological evaluation.

### Tissue Processing for single-cell RNA capture

After removal from the uterus, endometrial biopsies were immediately processed into single-cell suspensions and cryopreserved. Briefly, each endometrial biopsy was washed twice in PBS, cut into small pieces with a scalpel, and digested in 5 ml of IMDM (Gibco, 31980022) supplemented with 1 mg/ml Collagenase type I-A (Sigma-Aldrich, C2674) and 1 mg/ml Collagenase type II (Worthington, LS004174) at 37°C for 30-60 minutes. During digestion, the tissue-containing tube was inverted several times every 10 minutes until tissue pieces appeared visually dissociated. Enzymatic dissociation was terminated by the addition of 5 ml 4°C PBS. Cells were mechanically dissociated in a gentleMACS™ C Tube (Miltenyi Biotec, 130-093-237) using the gentleMACS™ Dissociator (Miltenyi Biotec, 130-093-235), with the Program mLung 02.01. The resulting cell suspension was filtered through a 100µm and 40µm strainer (Corning, 352340 and 352360) and rinsed with an additional 30 ml 4°C PBS. To remove red blood cells, cell suspension was pelleted by centrifugation (300g, 10 min), suspended in 6 ml of 1x red cell lysis buffer (Invitrogen, 00-4300-54), and incubated for 5 min at room temperature. The lysis was stopped by the addition of 35 ml 4°C PBS and endometrial cells were filtered using a 20µm strainer (pluriSelect, 43-50020-01). Following centrifugation (300g, 10 min), the cells were cryopreserved in 90% FCS (Sigma-Aldrich, F7524) 10% DMSO (Sigma-Aldrich, D8418), frozen in a Bicell freezing vessel (Nihon Freezer, 1-6263-01) at -80°C and subsequently stored long term in liquid nitrogen. For this project, 330 pipelles from 330 different donors were dissociated to single-cells and cryopreserved between 5/2020-12/2021.

### Single-cell RNA capture with 10x Genomics and sequencing with Illumina

Samples for 10x capture were selected by a thorough review of clinical data, pathology reports, menstrual cycle information, pipelle scores and single-cell quality. Selected endometrial single-cell aliquots were thawed in a 37°C water bath and suspended in 37°C IMDM supplemented with 10% FCS. After centrifugation (300g, 5 min), cell suspension was washed with 4°C PBS + 0.2% BSA (Sigma, A4503). After centrifugation (300g, 5 min), the cell pellet was carefully resuspended in 4°C PBS + 0.2% BSA to a concentration of ∼1*10^6 cells/ml and strained using a 35µm nylon mesh (Corning, 352235) or a 40µm Flowmi cell strainer (Sigma-Aldrich, BAH136800040) to ensure the absence of cell aggregates. Cell concentration and viability (propidium iodide-negative population) were measured after 5 min incubation of cell aliquot in 2 µg/ml propidium iodide (Sigma, 81845) using the portable flow cytometer Moxi Flow (ORFLO Technologies, cassette Typ MF-M, MXC021). Only samples with a post-thawing viability of >70% were included for single-cell capture.

Single-cell capture and cDNA library generation were performed with the 10x Genomics Chromium Single Cell Gene Expression workflow (Chromium NextGEM Single Cell 5’ Library and Gel Bead Kit v1.1, Chromium Controller). 25000 viable cells per sample were captured in a single reaction. cDNA libraries were balanced by shallow iSeq and sequenced with a NovaSeq 6000 System (Illumina, S4, 2 x 100 cycles).

### scRNA-seq data processing and atlas generation

The workflow was embedded in the workflow management engine Snakemake^75^ for automation and to ensure reproducibility. The software MultiQC^76^ was used for the quality control of the raw reads. Gene indexing, cell debarcoding, deduplication, read mapping, and estimation of transcript-level expression by pseudo-alignment were performed with the Salmon software package AlevinQC^77^. The Seurat^24^ and scater^78^ packages were used to perform quality control and visualization of the data on the sample-, cell- and gene-level. Included are the detection and removal of outlier cells based on transcript and gene metrics, detection of possible doublet cells, and batch effects. Samples not fulfilling the following quality criteria were excluded: median cell-wise mitochondrial expression < 30%, median number of genes per cell > 1000, and median number of unique transcripts per cell > 1000. Dimension reduction was done by selecting the 3000 highly variable genes that account for the most variation in a cell population.

### Principal component analysis

Sample-wise Principal Component Analysis (PCA) was performed on the sample-wise aggregated integrated transcripts^24^ with PCAtools.

### Menstrual cycle phase-specific marker genes

Mesenchymal or epithelial cells were subsetted from the entire single-cell atlas and samples 65 and 105 were excluded, due to their possible transitional state between periovulatory and secretory menstrual cycle phases. Menstrual cycle phase-specific markers were identified with the FindAllMarkers function from Seurat and filtered for adjusted p-value < 0.05 and avg_log2FC > 2.

### Cell-type identification

Cells were annotated with Symphony using Tan. et al. as reference. First, the entire dataset was annotated with the coarse annotation from Tan et al.^16^. Any annotations with a probability below 0.5 were discarded. Second, the entire dataset was split into the main cell types (mesenchymal, epithelial, endothelial, myeloid, and lymphoid) and subsequently annotated with the refined annotation from Tan et al.^16^. Any annotations with a probability below 0.5 were discarded. The marker genes of the cell types were confirmed with DotPlot^24^. Graph-based clustering with Louvain algorithm with multilevel refinement unveiled endometrial fibroblasts, C7 fibroblasts, and mesenchymal mural cell clusters specific to the menstrual cycle phase.

### Time trajectory of the cycle phase with monocle3

Time trajectory analysis was performed on the re-integrated epithelial cells with exclusion of ciliated, mesothelial, and MUC5B+ cells. Trajectories were deduced using the learn_graph function in the Monocle 3 v3_1.3.4 R package^79–81^, with a geodesic_distance_ratio set to 0.27. The root of the trajectory was determined by identifying the graph node close to cells from patients with the early menstrual cycle days 5, 6 and 7. Subsequently, cells were ordered along pseudotime using Monocle 3.

### Menstrual cycle phase evaluation

The menstrual cycle phase of each patient was assessed from serum progesterone levels, histological endometrium evaluation (if available) by two independent pathologists, and the transcriptional profile. Samples from women without measurable serum progesterone (< 2nmol/l) and menstrual cycle day ≤ 15 were considered to be in the proliferative menstrual cycle phase. Samples from women with measurable serum progesterone (> 2nmol/l), that clustered together with proliferative samples in PC1/PC2 and the UMAPs of all cells and epithelial cells were considered as periovulatory, samples that clustered separately as secretory. From the transcriptional profile, proliferative and periovulatory samples could not be distinctly identified. The evaluation of secretory into early-, mid- and late-secretory was performed under the consideration of the pathologist’s evaluation (if available) and the transcriptomic profile. Samples with the majority of stromal cells being in the dS3 cell cluster (separate cluster in UMAP of all cells) were considered as early- and mid-secretory, with a clear separation between early- and mid-secretory cells in the UMAPs of all cells or epithelial cells. The late-secretory stromal cells clustered completely separately in the UMAP of all cells with high frequencies of dS4 cells. This separation corresponds to the phase marker genes described in Wang et al. 2020. Marker genes for all 5 main cell types and each menstrual cycle phase are provided in Fig. 2d and Supplementary Table 2.

### Differential gene expression and pathway enrichment analysis

Cluster-wise differential expression analysis, comparing non-ENDO to mild-, severe- or all-ENDO, was performed with Muscat by fitting cell-level models on the RNA assay^27^ with 10x capture batch as covariate. Samples from the proliferative phase strict exclusion criteria were used as input (Supplementary Table 1). Only clusters were considered for testing with at least 3 ENDO and 3 non-ENDO samples with at least 50 cells per sample. Clusters too small for DEG testing were, if possible, aggregated with biologically similar clusters (e.g., pNK, NK1, NK2, NK3 to NK, Supplementary Table 3). The cut-off for differential expression was set at a p-value local adjusted of smaller than 0.05 and average expression > 18.53202 (inflection point). Ribosomal genes were excluded. Pathway enrichment analysis was performed with gProfiler^82^.

### Ligand-receptor interaction analysis

Ligand-receptor analysis was performed on the proliferative samples with strict exclusion criteria (Supplementary Table 1) using the R package CellChat with standard settings^53^.

### Cell cycle phase

Cell-cycle scores were assigned to mesenchymal cells with the Cell-Cycle vignette from Seurat ^24^, based on the G2/M and S phase markers genes^83^. Two mesenchymal cell clusters (Seurat clusters 2 and 4) contained cells with G2/M and S cell-cycle scores above 1 and were considered as cycling mesenchymal cells.

### ScaiVision analysis

Endometriosis predictions were carried out using the interpretable neural network algorithm CellCnn^65^, implemented in PyTorch in the ScaiVision platform (version 1.6.3, Scailyte AG), similar to Roussel et al.^84^. Briefly, this is a supervised machine learning algorithm that trains a convolutional neural network with a single hidden layer to predict sample-level labels using single-cell data as inputs.

The scRNA data of the proliferative, no progesterone samples with more than 5000 cells per sample (Supplementary Table 1) were included in the ScaiVision network training, using either all cells or only cycling fibroblast cells^24,83^. Each sample was given a label corresponding to the Endometriosis status of the patient from which the endometrial biopsy was taken (Endometriosis or Control). For model training, we used a nested cross-validation scheme. In the outer loop, we followed a 5-fold Monte Carlo cross-validation (MCCV) scheme in which ∼20% of samples were set aside for evaluation for each fold, resulting in 5 independent evaluation datasets. The remaining samples (∼80%) of each fold were further allocated to 5 MCCV splits of training (70%) and validation (30%) in a stratified manner to maintain the relative proportions of each class (technical batches and endpoints non-ENDO/ENDO), resulting in 25 training and validation datasets. This inner loop was used for hyperparameter tuning and selection.

Four different feature selection methods were used to reduce the dimensionality of the data prior to model training: 1) PCA^67^, 2) CorrgPCA 3) differentially expressed genes (“diffgen”)^68^, and 4) highly variable genes (“hvg”)^85^. CorrgPCA, Scailyte’s proprietary dimension reduction technique, first groups the highest variable genes^85^ based on pairwise correlation. It then computes a principal component analysis on those gene groups separately and selects the 200 components capturing the highest amount of variance.

Per training dataset and feature selection method, 50 ScaiVision models with varying hyperparameters were trained on the dimension-reduced training data. To generate input data for training ScaiVision, sub-samples of 200 cells, termed multi-cell inputs (MCIs), were chosen randomly from each sample independently. For each training epoch, 1000 MCIs from each label class (Endometriosis or Control) were presented to the network in random order. 50 independent networks were generated for each feature selection method using hyperparameters randomly chosen from the following options: i) number of filters: (3, 5, and 10), ii) top-k pooling percentage: (20), iii) dropout probability: (0.3, 0.4, and 0.6), iv) learning rate: (0.001, 0.003, and 0.01), and v) weight decay: (0.00001, 0.0001, 0.001, 0.01, and 0.1). Training was performed with a batch size of 64. Each network was trained for a maximum of 50 epochs, or until the validation loss no longer decreased for 20 consecutive epochs. At the end of the training, the weights from the epoch with the lowest validation loss were returned.

For each of the outer MCCV folds and each feature selection method, 3 trained models were selected which performed best on their respective validation dataset and evaluated by computing AUC scores for the independent evaluation dataset, yielding 3 best-performing models per feature selection method and fold. We continued our analysis with those feature selection methods for which the best selected models achieved an AUC ≥ 0.8 on at least 3 of the 5 folds.

Models deriving from different feature selection methods use different sets of input genes; additionally, each model has the potential to learn different weights for each input feature. In order to derive a unified gene signature for PCA and CorrgPCA, the top network was picked from each of the 5-folds. Only networks that had an accuracy of 0.66 or above and an AUC of 0.78 or above on the held-out evaluation set were picked. In the case of ties, the network with the lowest overlap in the response distributions between endometriosis and control predictions was picked. Folds that did not have a single network satisfying these criteria were excluded from further analysis.

Next, for each of the networks, importance scores were calculated, using the deepLIFT^86^ and integrated gradients methods implemented in the Captum software^87^, providing a metric for the importance of each feature for driving the prediction of either endometriosis or control. Integrated gradients importance scores were averaged feature-wise across the deepLIFT and integrated methods and across all the samples within a group (endometriosis or control), and the absolute difference between the integrated gradients importance scores for endometriosis prediction and that for control prediction was calculated for each feature. Next, the top 20% of features, ranked by the mean absolute difference of integrated gradients importance scores, were selected for further consideration. For corrGPCA, each of the features (CorrgPCs) was mapped back to the top 20% of genes associated with the feature (as ranked by absolute PC loading of the gene). If the feature had fewer than 10 genes associated with it, then the top 2 genes were picked, and for singleton gene sets, the single gene was picked. Features associated with PCA were treated analogously, however, due to the non-sparse nature of PCs relative to corrGPCs, only the top 1% of genes were picked per integrated gradients importance score-ranked PC per feature.

To identify cells driving the prediction of our models, cells targeted by the models selected during the evaluation were examined. Each of the models contains different filters, which collectively form the model’s prediction. The combination of weights from each filter with the expression measurements through a weighted average gives a score per filter. This score represents how strongly the cells respond to the filter. To pinpoint predictive cells throughout folds, we examined all the filters and called a cell type as predictive if it was consistently selected in different folds.

### Statistics and Reproducibility

No formal sample-size calculation was performed. From 330 participants with cryopreserved endometrial biopsies, we included most meeting our stringent criteria. All data exclusion criteria were pre-established and data was only excluded if it did not meet the quality criteria, as described in detail above. For cell type frequency, differential gene expression and ligand-receptor analysis stringent patient exclusion criteria were applied (Table 1), due to a potential influence of these clinical parameters. To compare cell type frequencies across menstrual phases, we utilized 23 samples from the proliferative phase, 18 from the periovulatory phase, and 13 from the secretory phase, adhering to stringent clinical exclusion criteria. Retrospective alignment with a menstrual cycle atlas showed perfect phase correspondence. For differential gene expression and ligand-receptor analysis, the 12 samples from women with endometriosis and 11 without endometriosis from the transcriptionally homogeneous proliferative menstrual cycle phase were used, with stringent clinical exclusion criteria applied, ensuring reliable results.

## Data availability

The raw data and processed Seurat object are available on the European Genome-phenome Archive (EGA) with controlled access (accession number pending) and may be shared upon request. The data are not publicly available because they contain sensitive information that could compromise the privacy of research participants.

## Code availability

Code used for data analysis in this study is available at: https://github.com/DuempelmannLea/endometriosis_endometrium_scRNA_atlas.

## Acknowledgements

The authors thank all participants for their valuable participation in the study and their sample donation, the physicians, nurses, and study nurses involved in patient recruitment and sample collection, and Anne Vaucher for processing biological samples. The authors further thank the Next Generation Sequencing (NGS) Platform (Pamela Nicholson and Daniela Steiner) for scRNA library preparation and sequencing, the Steroid Laboratory (Michael Grössl and Clarissa Vögel, Nephrology depart., Inselspital, Bern) for the progesterone measurements, the Translational Research Unit (TRU) at Institute Of Tissue Medicine and Pathology (Irene Centeno Ramos, Paulina Brönnimann) for patient sample provision and H&E stainings and the Interfaculty Bioinformatics Unit (Heidi Tschanz-Lischer) for time trajectory analysis.

## Author contributions

M.D.M. and P.N. conceived and supervised the project and secured funding. L.D. and S.S. designed, supervised, and performed experiments/analyses. J.S. and T.A. performed experiments. A.D., D.G., S.M., and R.L. performed data analysis. S.C. supervised data scientists. W.S. and H.B. evaluated the menstrual cycle stage based on H&E staining. Authors were involved in conceptual discussions. L.D. and B.M. wrote the manuscript and L.D. prepared the figures. All authors discussed results and commented on the manuscript.

## Additional information

Supplementary Information is available for this paper.

Correspondence and requests for materials should be addressed to L.D. or M.D.M..

## Additional information

Supplementary Information is available for this paper.

## Extended Data Figures and Legends

**Extended Data Fig. 1.**
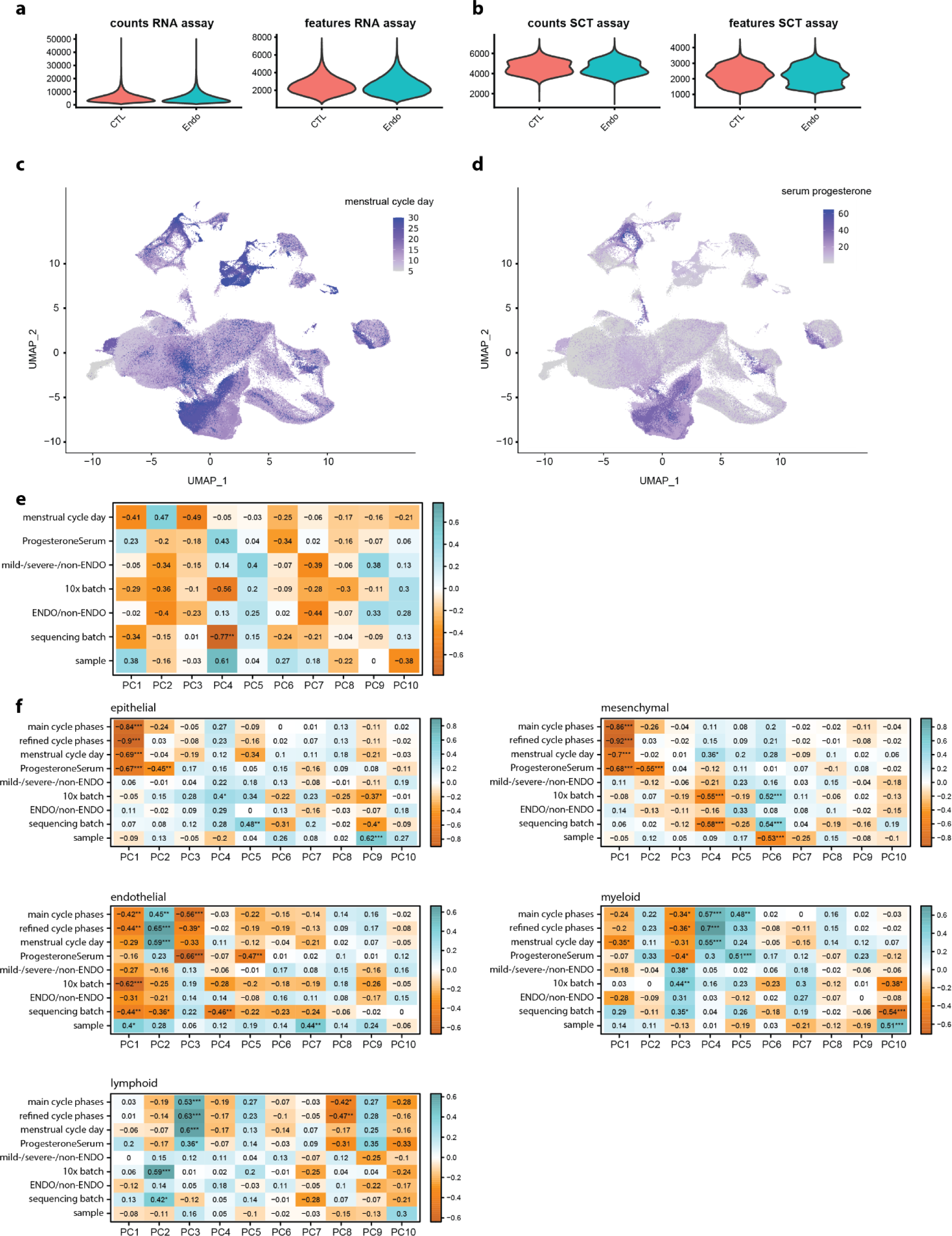
Additional parameters and features of the endometrium single-cell atlas. **(a,b)** Violin plots illustrating counts and features from RNA assay (a) and SCT assay (b), show no discernable differences between ENDO and non-ENDO. **(c,d)** UMAPs of the entire endometrium single-cell atlas, colored by menstrual cycle day (c) and progesterone in serum (d), show a clear visual separation of cells by menstrual cycle day and serum progesterone concentration. **(e)** Eigencorplot showing the effect of different co-variants of the atlas dataset on principal components (PCs) of the proliferative samples with strict exclusion criteria with Pearson correlation values and multiple-testing adjustment with the Benjamini-Hochberg procedure. No significant correlation was identified between menstrual cycle phase parameters or the endometriosis status and the initial ten PCs. Statistical significance symbols represent: *** <0.001, ** <0.01, * <0.05. **(f)** Eigencorplot showing the effect of different co-variants of the atlas dataset on principal components (PCs) of the main 5 cell types (epithelial, mesenchymal, endothelial, myeloid, lymphoid) with Pearson correlation values and multiple-testing adjustment with the Benjamini-Hochberg procedure. Menstrual cycle phase parameters significantly correlate with high PCs in all main cell types. In contrast, only in myeloids, a significant correlation was identified between the endometriosis status and the initial ten PCs of any main cell type. Statistical significance symbols represent: *** <0.001, ** <0.01, * <0.05.

**Extended Data Fig. 2.**
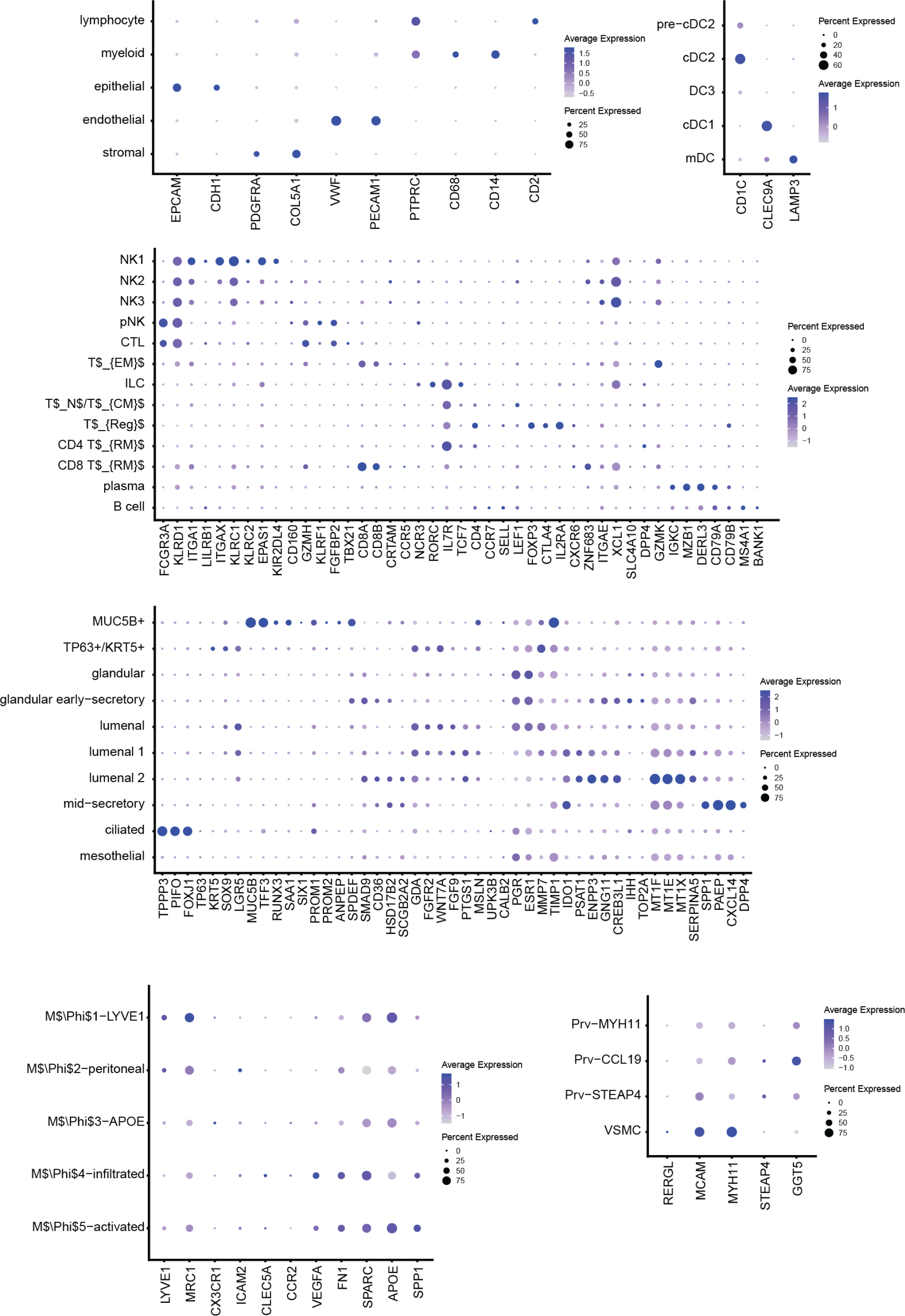
Marker gene expression from our dataset corresponds to the reference. Expression of the marker genes for the different cell types corresponds to the cell type-specific expression shown in Tan et al. 2022^16^.

**Extended Data Fig. 3.**
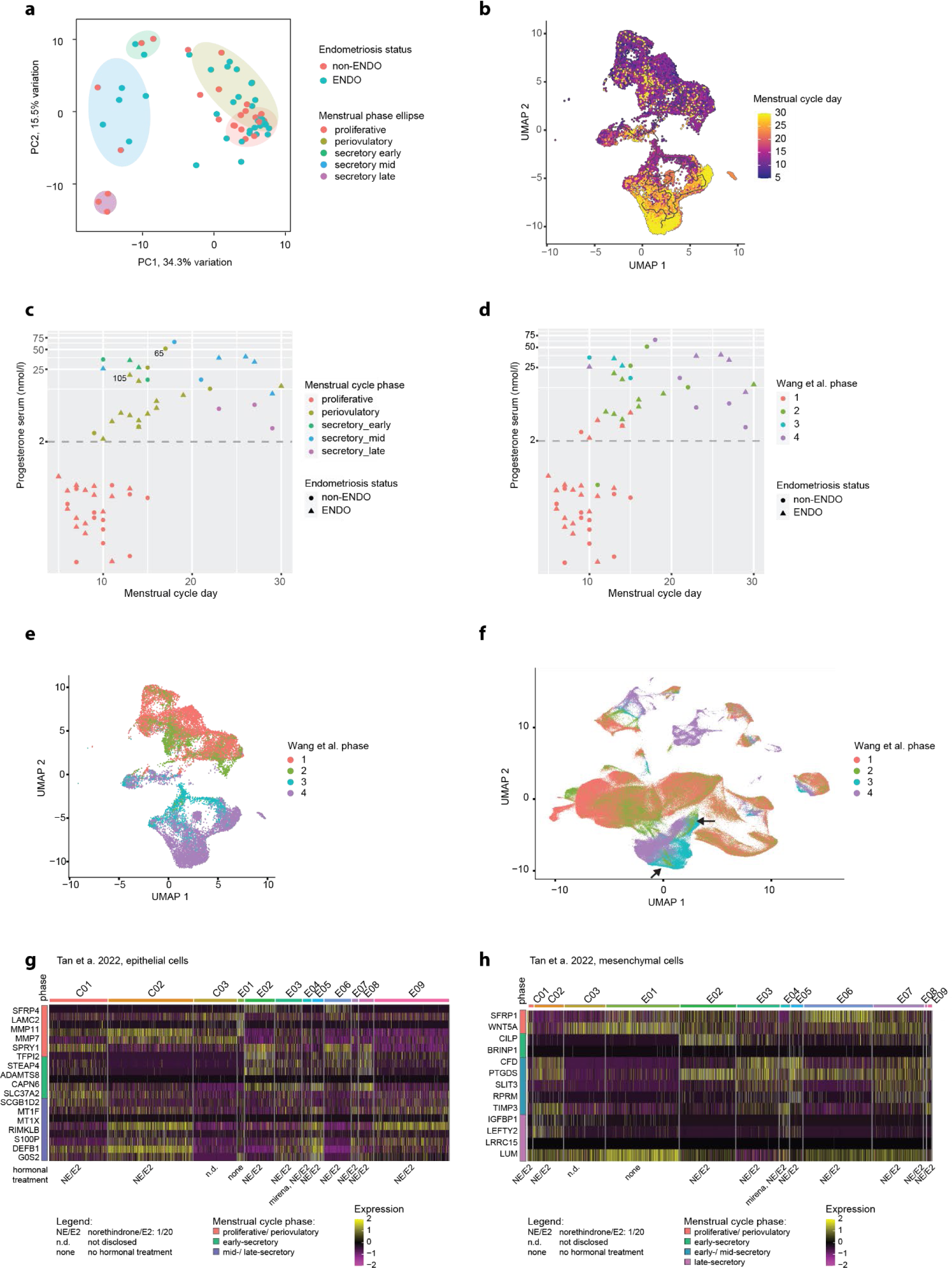
Strong correlation of menstrual cycle day and phase with Monocle pseudo time prediction and reference menstrual cycle single-cell atlas. **(a)** Scores plot of the first two principal components PC1 and P2 from sample-wise whole transcriptome PCA of the SCT assay, colored by endometriosis status. Ellipses mark the different menstrual cycle phases (same as in Fig. 2a). ENDO and non-ENDO do not cluster distinctively in any menstrual cycle phase. **(b)** UMAP of re-integrated epithelial cells colored by menstrual cycle day. A clear progression from early to late menstrual cycle day is visible. **(c,d)** Scatter plots showing the correlation of menstrual cycle day and serum progesterone with our menstrual cycle staging (c) and median Wang phases (d), transferred with Symphony. There is a perfect correspondence between our cycle phase evaluation and the Wang phases (proliferative phase with Wang phases 1 and 2, early-secretory phase with Wang phase 3, and mid-/late-secretory phase with Wang phase 4) **(e,f)** UMAPs of the entire dataset (left) and the re-integrated epithelial cells (right), colored by the Symphony transferred Wang phases, show a clear visual separation between phase 1/2, phase 3, and phase 4. The cells from phase 2 in the early-/mid secretory mesenchymal cluster (arrows) are from sample 65 and 105. **(g,h)** Heatmaps showing the expression of menstrual cycle phase markers (as in Figure 2d) of the eutopic endometrial epithelial cells (**g**) and mesenchymal cells (**h**) from the Tan et al. 2022 single-cell atlas. C01-C03 are control patients and E01-E09 endometriosis patients. The hormonal treatment of each donor is given below the heatmap. The menstrual cycle phase of the highest gene expression is annotated between gene names and heatmap.

**Extended Data Fig. 4.**
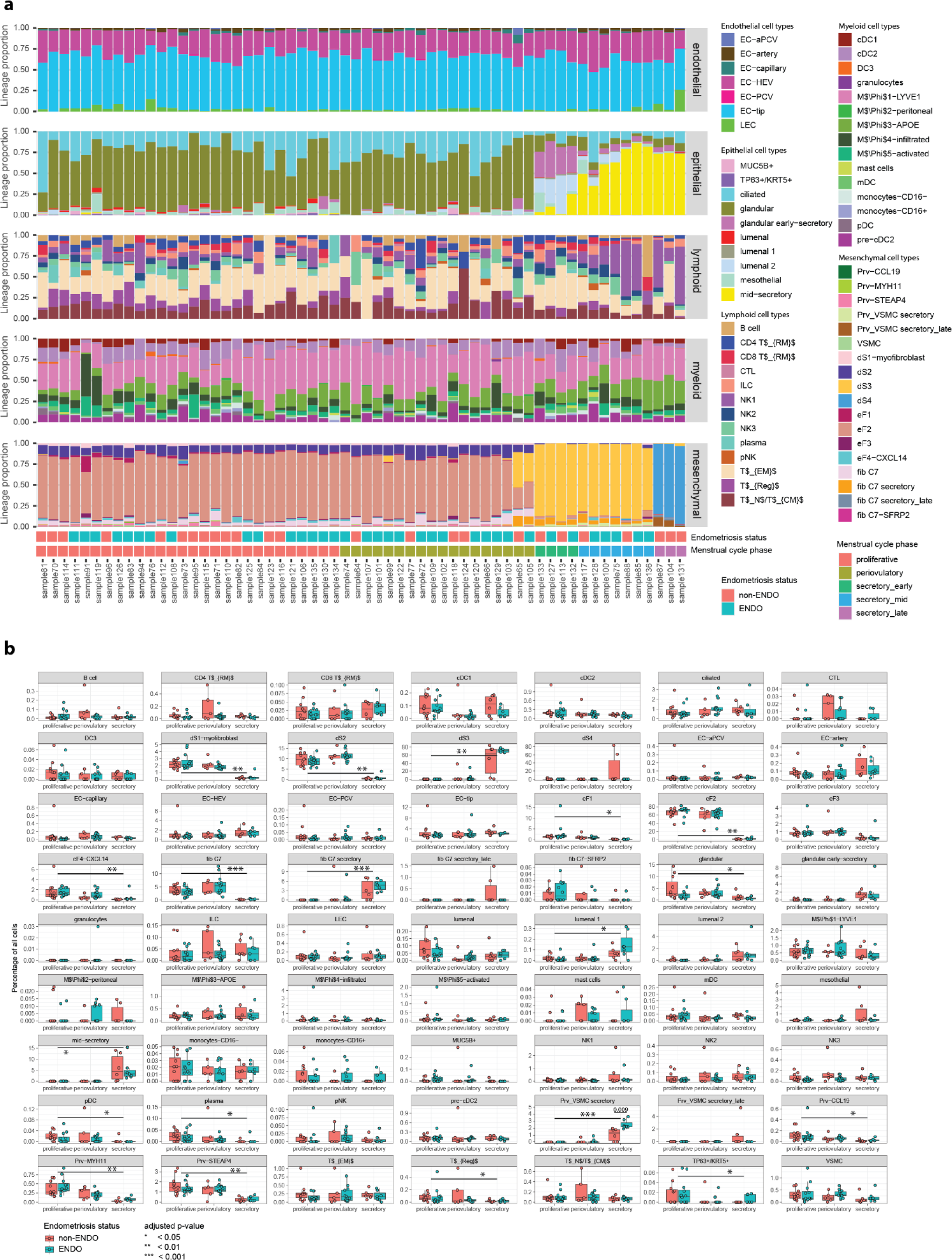
Cell type frequency changes are mainly driven by the menstrual cycle, not the endometriosis status. **(a)** Barplot of cell type frequencies per sample, sorted by menstrual cycle phase and pseudo time and divided by the main 5 cell types. There is no clear trend separating ENDO and non-ENDO. **(b)** Box plots of refined cell type frequencies between the main menstrual cycle phases. p-values for cell type frequency change between the proliferative and secretory menstrual cycle phase were calculated with a two-tailed Student’s t-test and adjusted with the Benjamini-Hochberg Procedure, * adjusted p-value < 0.05, ** adjusted p-value < 0.01, *** adjusted p-value < 0.001. Secretory mural cell (Prv_VSMC secretory) are 2.7-fold more abundant in ENDO than in non-ENDO (unadjusted p-value = 0.009).

**Extended Data Fig. 5.**
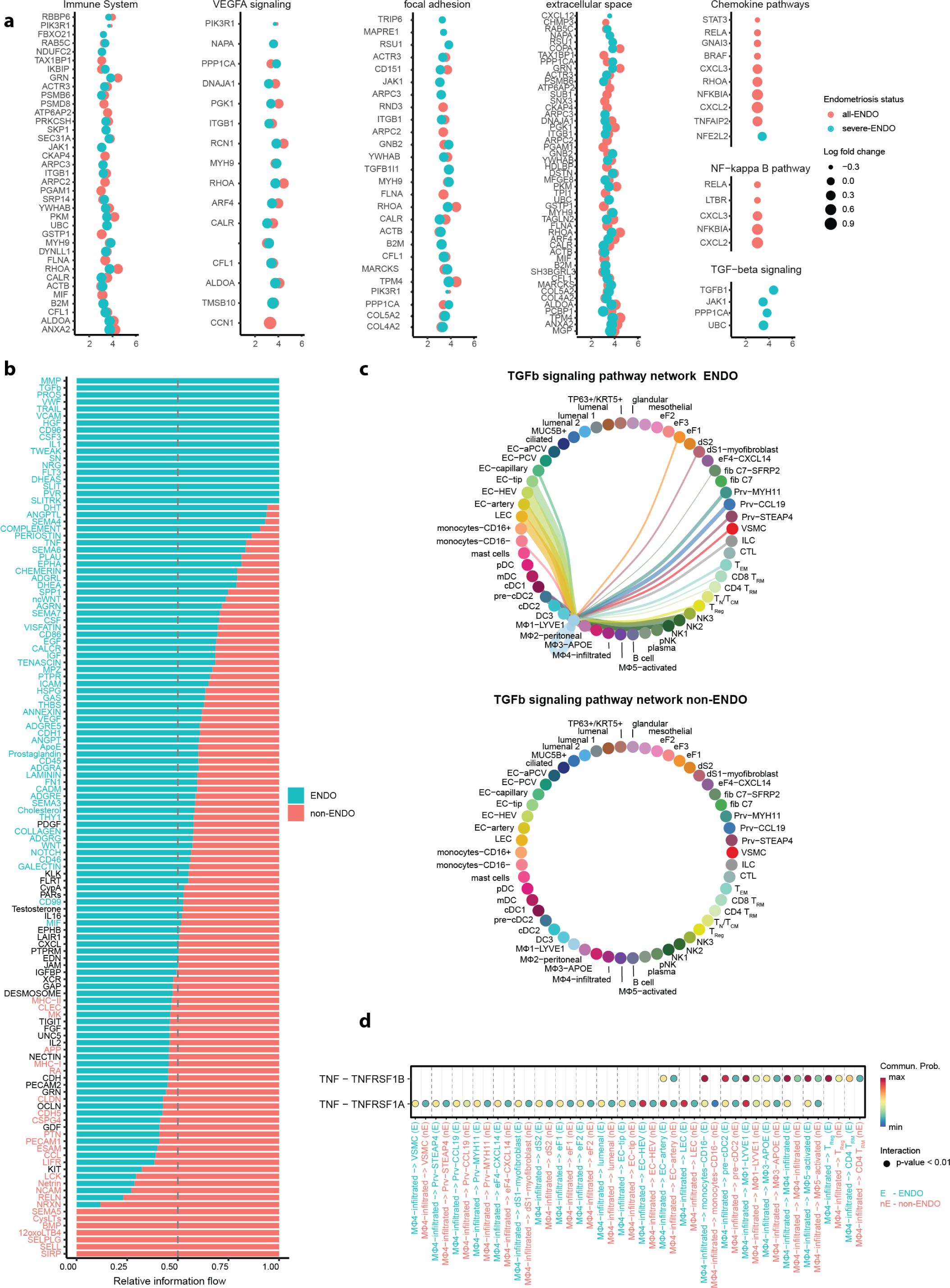
Inflammation, Adhesion, Proliferation, and Angiogenesis pathways and ligand-receptor interactions are upregulated in endometriosis endometrium. **(a)** Genes underlying selected pathways enriched in the cluster- and stage-wise functional enrichment analysis. **(b)** Signaling pathways with relative information flow from CellChat analysis, ranked by overall information flow within the inferred networks between ENDO and non-ENDO. Signaling pathways with a red font have overall enriched activity in ENDO and pathways with a turquoise font have overall enriched activity in non-ENDO. **(c)** Circle plot of TGFb signaling pathway between all cell types in ENDO (top) and non-ENDO (bottom, no significant interactions). **(d)** Bubble plot of selected communication with significant interaction probabilities (unadjusted p-value < 0.01) within the TNF signaling pathway. Further p-value adjustment is not recommended by the CellChat author.

**Extended Data Fig. 6.**
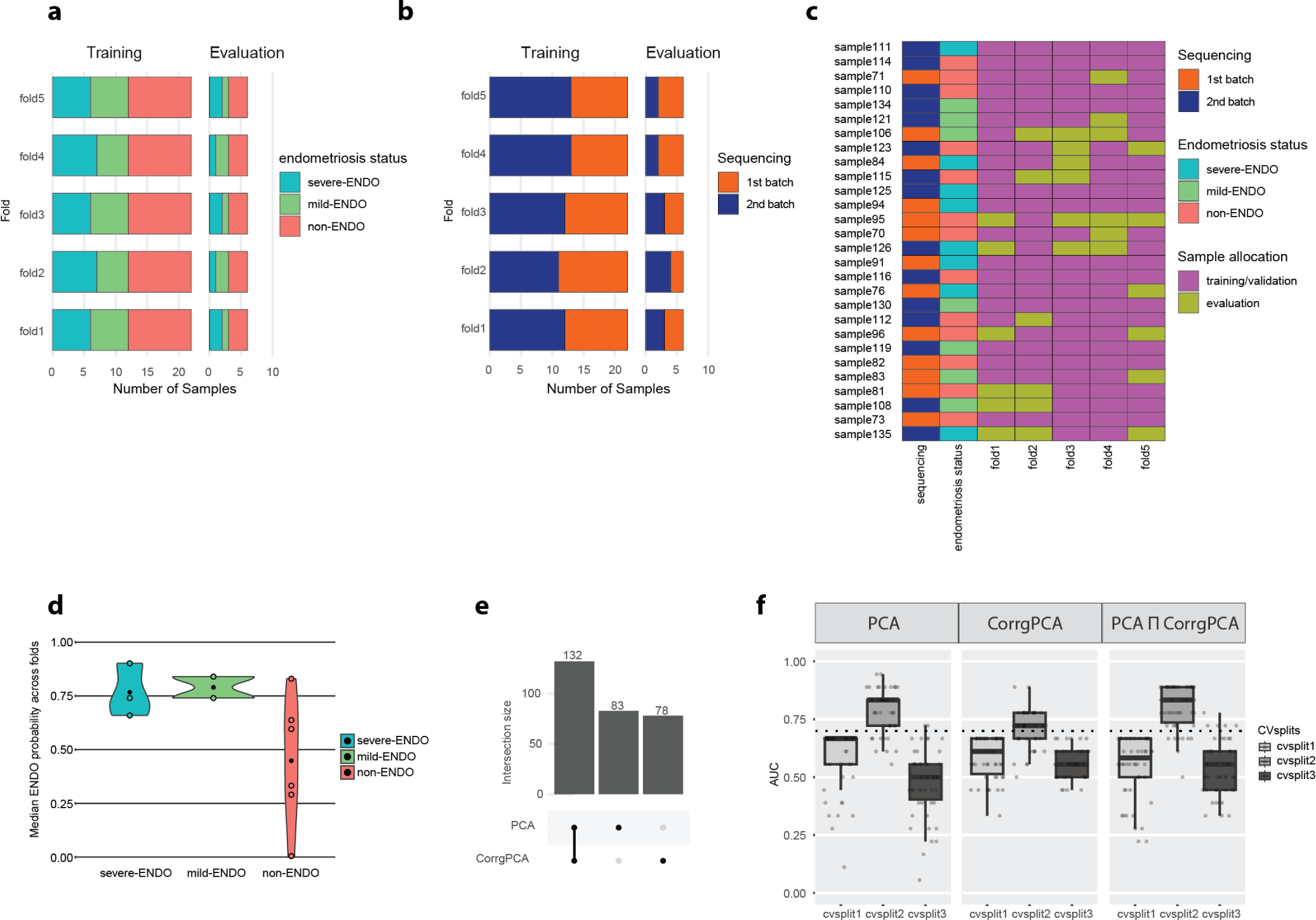
Parameter distributions and predictions of the ScaiNET model training. **(a,b)** Endometriosis status and sequencing runs are similarly distributed between the training, validation, and evaluation folds. **(c)** Heatmap showing the parameters of each sample, as well as its fold-wise allocation to training/validation and evaluation through the 5-fold Monte Carlo cross-validation scheme. **(d)** Exemplary violin plot of median endometriosis probability (black dot) from the top-ranked learner from the CorrgPCA 3-CV prediction for severe-ENDO, mild-ENDO, and non-ENDO samples, with no discernible difference between mild-ENDO and severe-ENDO prediction. **(e)** UpSet plot of the unified gene signature from PCA (n = 215) and CorrgPCA (n = 210) reveals a substantial 60% overlap (n = 132). **(f)** CellCnn prediction based on the unified PCA and CorrgPCA signature on 3-CV splits of the non-proliferative samples performs poorly with median AUCs below 0.7 (dotted line).

**Extended Data Fig. 7.**
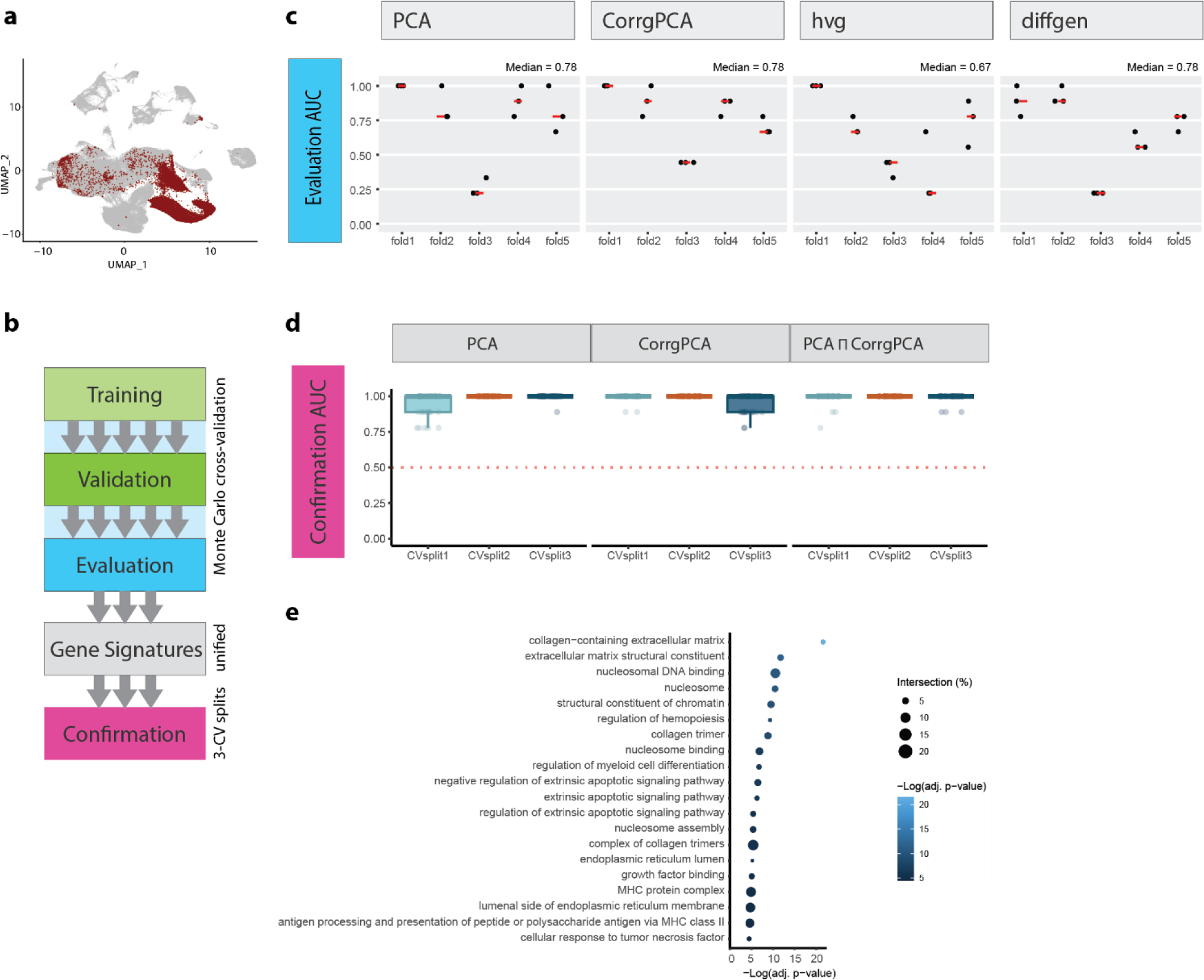
ScaiNET prediction of endometriosis based on cycling mesenchymal cells. **(a)** UMAP of endometrium single-cell atlas with cycling mesenchymal cells highlighted. **(b)** Workflow outlining the ScaiVision model training and gene signature confirmation process. **(c)** Boxplots of evaluation AUCs split by feature selection methods (PCA, CorrgPCA, hvg, and diffgen). The median evaluation AUC per fold is represented by the dark red line. The median AUC per plot is shown on its top right. PCA, CorrgPCA and diffgen performed best in the evaluation with median AUCs of 0.78. **(d)** Boxplot of the confirmation AUC for the unified PCA and CorrgPCA gene signatures and their intersect (PCA ∩ CorrgPCA), showcasing median AUCs of 1 for each CV split. **(e)** Top 20 terms and pathways, ranked by the adjusted p-value, highlight structural terms, cell differentiation, and responses to diverse stimuli.

## Supplementary Tables

**Supplementary Table 1 | Clinical data of study participants and sample details.**

**Supplementary Table 2 | Menstrual cycle specific marker genes of the main five cell types.**

**Supplementary Table 3 | Cell type frequency stratified by sample, endometriosis status, and menstrual cycle stage.**

**Supplementary Table 4 | Cluster and stage-wise differentially expressed genes of the proliferative menstrual cycle phase samples with stringent exclusion criteria applied.**

**Supplementary Table 5 | Pathways and terms revealed by cluster- and stage-wise gene enrichment analysis.**

**Supplementary Table 6 | ScaiNET PCA and corrgPCA gene signatures.**

**Supplementary Table 7 | Cell-type-wise percentage of cells within the top 10% of filter scores for ENDO or non-ENDO prediction.**

